# An atlas of functional ensembles in the zebrafish hindbrain links structural and molecular features to behavior

**DOI:** 10.64898/2026.09.30.752102

**Authors:** Sophie Aimon, Heike Naumann, Armin Bahl

## Abstract

The vertebrate hindbrain sustains functions essential for survival and behavioral control, yet how it operates as a whole remains poorly understood. Here we map its functional organization in larval zebrafish, whose transparency permits imaging of the entire hindbrain. We first curate established functional populations from the literature into an atlas annotated with putative mammalian counterparts. We then complement this library with ensembles of co-active neurons extracted without prior labels from whole-hindbrain light-field recordings in behaving larvae. We identify 70 ensembles that recur across individuals and recover most of them in independent light-sheet datasets. Ensembles also align with spatial gene-expression patterns and share their bilateral symmetry. Behavioral characterization identifies known eye- and tail-movement-related populations and reveals new associations with fin and mouth movements. Combining co-activity, molecular signatures, and neuronal projections suggests a candidate recurrent inter-regional circuit. Our atlas thus provides a common spatial reference for linking functional, molecular, and anatomical data to guide circuit discovery.

**HIGHLIGHTS:**

- hindbrain ensembles recur across fish, and most are robustly detected across methods
- ensembles match gene expression and share its bilateral symmetry
- track movement of multiple body parts around swimming onset
- maps link activity, molecular and projection data to guide circuit discovery

## INTRODUCTION

The vertebrate hindbrain sits at the base of the neuraxis and sustains functions essential for survival (Schröder et al., 2023; Striedter et al., 2020), including posture and gaze stabilization (Straka and Baker, 2013), locomotion (El Manira, 2026), and arousal and visceral control (Zhao et al., 2026), while also contributing to broader brainstem networks for orienting and escape (Arber and Costa, 2022; Branco and Redgrave, 2020). Yet how its neuronal populations are organized into a coordinated functional system remains poorly understood. Its component nuclei have been characterized in great anatomical, molecular, and physiological detail, but recordings restricted to individual nuclei cannot directly reveal how activity is temporally organized across the hindbrain. Broad optical access is difficult in mice because the brainstem lies beneath the cerebellum; implanted lenses or prisms trade field of view and speed for depth (Agetsuma et al., 2025). Electrode arrays can sample brainstem activity directly, but each insertion covers only a limited volume, so brain-wide mammalian maps are assembled across many recordings and animals rather than observed simultaneously in one (Meshulam et al., 2025). In humans, ultra-high-field imaging can cover the whole structure at once, resolving 58 brainstem nuclei and their cortical connectivity (Hansen et al., 2024). However, these measurements operate at hemodynamic timescales and use predefined anatomical parcels, not populations identified from activity. How hindbrain nuclei act together as a coordinated system therefore remains far less well-described than their individual properties.

A useful entry point for describing such distributed organization is the neuronal ensemble. Across nervous systems, coordinated groups of co-active neurons have been described as cell assemblies, neuronal ensembles, or functional modules, and are increasingly treated as a fundamental unit of neural computation, complementary to descriptions based on single neurons or anatomically defined brain regions (Buzsáki, 2010; Harris, 2005; Hebb, 1949; Saxena and Cunningham, 2019; Yuste et al., 2024). This framework has been applied across species and brain regions, from mammalian cortex (Miller et al., 2014) to whole-brain recordings in small model animals (Aimon et al., 2019; Chen et al., 2018; Dupre and Yuste, 2017). Its power is that ensembles are defined by what neurons do together, rather than by where they sit or which marker they express. Applied systematically to a vertebrate hindbrain, this approach could bring together an extensive body of region-by-region physiology into a unified, activity-defined map.

The small size and transparency of larval zebrafish make it possible to examine the organization of neuronal activity across the entire hindbrain within individual animals. Extensive previous work has identified neuronal populations involved in gaze stabilization and vestibular control (Beiza-Canelo et al., 2023; Migault et al., 2026; Miri et al., 2011; Schoppik et al., 2017), sound processing (Poulsen et al., 2021), locomotion and turning (Carbo-Tano et al., 2023; Dunn et al., 2016; Kimura et al., 2013; Lau et al., 2025), escape (Marquart et al., 2019), orienting and heading representation (Petrucco et al., 2023), and sensorimotor transformation (Bahl and Engert, 2020; Dragomir et al., 2020; Slangewal et al., 2026) (see also **Supplementary Table 1**). These studies have revealed many of the building blocks of hindbrain function. Compiling these populations and their functional annotations in a common spatial reference would provide a basis for comparing activity-defined ensembles with established functional systems and identifying ensembles without a clear counterpart among the compiled populations. Annotating the compiled populations with proposed mammalian counterparts would consolidate comparative evidence dispersed across specialized literatures and link this zebrafish atlas to corresponding populations in the mammalian brainstem and cerebellum.

Beyond individual circuits, brain-wide studies in larval zebrafish have revealed reproducible functional organization without predefined anatomical regions. Response clustering identified convergent sensorimotor transformations (Chen et al., 2018), while a compositional restricted Boltzmann machine recovered spatially conserved neural assemblies whose organization reflected structural connectivity (Van Der Plas et al., 2023). Correlational networks likewise followed anatomical connectivity (Légaré et al., 2025), and immunostaining for the activity marker pERK mapped brain-wide responses across stimuli and behaviors (Randlett et al., 2015). Yet each study defined functional organization within its own experimental and analytical framework, rather than testing a common set of units across datasets. A common set of co-active spatial units recurring across individuals, methods, and imaging modalities would allow direct comparison across studies. Registered to a common reference, these units could also be compared with gene-expression patterns and anatomical parcels, testing whether co-activity follows molecular organization (Kunst et al., 2019; Marquez-Legorreta et al., 2026; Shainer et al., 2023).

The hindbrain’s bilateral organization provides a particularly clear case in which to examine this correspondence. Many identified motor and premotor neurons and populations have recognizable counterparts across the midline (Collins et al., 2025; Hildebrand et al., 2017; Metcalfe et al., 1986). In well-studied hindbrain systems, including vestibular and escape circuits, this scaffold is embedded in conserved developmental patterning and supports functions ranging from bilateral coordination to opponent interactions that can resolve left–right action choices (Koyama et al., 2016; Straka and Baker, 2013). Anatomical pairing, however, does not ensure that corresponding left and right populations are co-active. Left and right populations may operate largely independently, contribute jointly to a single bilaterally symmetric ensemble, remain distinct but coactive as unilateral partners, or engage in opponent dynamics (Aksay et al., 2007; Wolf et al., 2017). To date, studies of these bilateral activity motifs have focused mainly on individual, predefined populations, leaving their prevalence and spatial distribution across the hindbrain incompletely characterized.

Interpreting these ensembles also requires behavior to be measured broadly enough to distinguish the actions that the hindbrain coordinates. Whole-brain imaging studies in larval zebrafish have focused mainly on swimming and eye movements. Pectoral-fin and orofacial movements have been characterized behaviorally and in targeted circuit studies (Green et al., 2011; McArthur et al., 2023; Thorsen et al., 2004; Uemura et al., 2020), but have not been related to brain-wide neural activity. Because these movements can accompany swimming (Bianco et al., 2011; Green et al., 2011), neural activity correlated with swimming may partly reflect accompanying fin or orofacial movements. Simultaneous, bout-resolved tracking of all four effectors is therefore needed to determine how hindbrain ensembles coordinate movements across body parts and to disambiguate their behavioral associations. Relative timing of activity and behavior provides a complementary axis for interpreting ensemble function. Anticipatory, concurrent, and lagging activity offer clues to roles in sensory processing, motor decision-making, movement execution, and sensory reafference. Finally, ensembles that recur reliably across individuals but remain weakly associated with this expanded set of sensory and behavioral variables may encode unmeasured sensory dimensions, actions, or internal states. Their spatial maps provide starting points for investigating the underlying circuits.

Here we combine a comprehensive literature-based atlas of established hindbrain functional populations with unsupervised analysis of whole-hindbrain light-field imaging in behaving larval zebrafish. First, we compile established zebrafish hindbrain populations into a common spatial reference and annotate their putative mammalian counterparts where comparative evidence is available. We then extract co-active ensembles from whole-hindbrain activity and show that many recur reproducibly across individuals, with most also robustly recovered in independently acquired datasets using alternative decomposition algorithms. By applying the same unsupervised decomposition strategy to a registered gene-expression atlas, we find that functional ensembles show greater spatial correspondence to expression-derived components than to classical anatomical parcellations. We further show that these ensembles are strongly bilaterally organized, in a manner that matches the symmetry of hindbrain gene expression. We also track eye, tail, fin, and mouth movements simultaneously and find that many ensembles are broadly coupled to locomotor bouts, while more selective associations with individual behaviors are confined to limited subsets of ensembles, including a separable orofacial channel active in the absence of tail movement. Across the atlas, responses around swim onset span a graded, stereotyped range of latencies, from anticipatory to lagging. Finally, we combine co-activity, molecular signatures, and neuronal projections to propose a candidate recurrent circuit, illustrating how ensembles weakly explained by the recorded variables can guide circuit discovery. Together, these results define an activity-based atlas of a vertebrate hindbrain and provide a framework for relating multi-region circuit organization to anatomy, molecular identity and behavior.

## RESULTS

### A curated atlas of established zebrafish hindbrain functional populations

To provide a literature-derived reference for interpreting the activity-defined ensembles, we assembled established larval zebrafish hindbrain neuronal populations in a common anatomical reference (**Fig. 1**). This reference allowed us both to assess whether unsupervised whole-hindbrain analysis recovered established functional systems and to identify reproducible ensembles outside current functional maps. We also annotated putative mammalian counterparts where comparative evidence was available, providing a comparative extension of the zebrafish reference (**Fig. 1** and **S1**, **Supplementary Table 1**).

**Figure 1.**
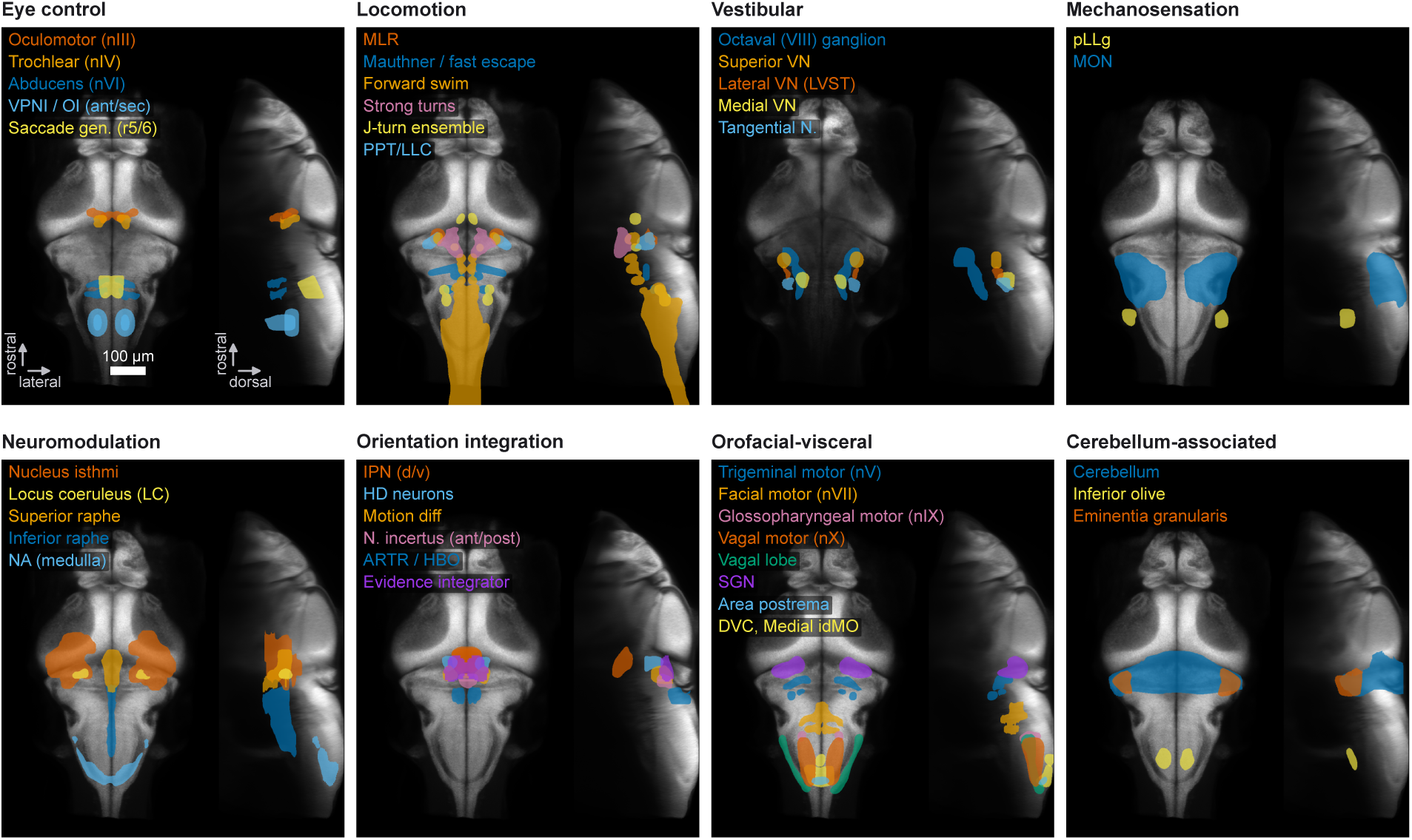
A literature-based atlas of established functional populations in the larval zebrafish hindbrain. Established functional regions and populations compiled from the literature and displayed as colored masks over the mapzebrain reference brain in grayscale. The grayscale background is a mean projection of reference-brain slices that intersect the displayed masks. Each functional category is shown in dorsal (left) and lateral (right) views. Populations are organized into eight non-exclusive groups based on functional roles or membership in established neural systems: eye control, locomotion, vestibular, mechanosensation, neuromodulation, orientation integration, orofacial–visceral, and cerebellum-associated. Masks were taken from existing atlas regions where available and otherwise drawn from published figures and descriptions. **Supplementary Table 1** provides the functional annotations, mask sources, and construction details. Proposed mammalian counterparts are summarized in related **Fig. S1**. **Abbreviations: Eye control:** nIII, oculomotor nucleus; nIV, trochlear nucleus; nVI, abducens nucleus; OI, oculomotor integrator; VPNI, velocity-to-position neural integrator; Saccade gen., saccade generator. **Locomotion:** LLC, long-latency C-start; MLR, mesencephalic locomotor region; PPT, prepontine nucleus. **Vestibular:** VN, vestibular nucleus; LVST, lateral vestibulospinal tract. **Mechanosensation:** MON, medial octavolateralis nucleus; pLLg, posterior lateral-line ganglion. **Neuromodulation:** LC, locus coeruleus; NA, noradrenergic/catecholaminergic groups. **Orientation integration:** IPN, interpeduncular nucleus; ARTR, anterior rhombencephalic turning region; HBO, hindbrain oscillator; HD, heading-direction neurons; Motion diff, motion-onset/direction-selective neurons. **Orofacial– visceral:** DVC, dorsal vagal complex; idMO, intermediate dorsal medulla oblongata; nV, trigeminal motor nucleus; nVII, facial motor nucleus; nIX, glossopharyngeal motor nucleus; nX, vagal motor nucleus; SGN, secondary gustatory nucleus. **Other:** ant/post, anterior/posterior; d/v, dorsal/ventral; N., nucleus; r5/6, rhombomeres 5 and 6.

We compiled a literature-based reference of hindbrain functional organization by surveying published studies for functional regions and populations, defined here as groups of neurons that are strongly co-activated or share a common function. When a population corresponded to an existing atlas parcel, we used the relevant mapzebrain (Kunst et al., 2019) or Z-Brain (Randlett et al., 2015) mask. For populations not delineated in either atlas, or occupying only part of an existing parcel, we generated an operational mask or subregion in atlas space from published figures, descriptions, registered functional maps, or expression maps (**Supplementary Table 1**) (Kunst et al., 2019; Randlett et al., 2015; Shainer et al., 2023).

We organized the masks into eight non-exclusive functional groups: eye control, locomotion, vestibular, mechanosensation, neuromodulation, orientation integration, orofacial-visceral, and cerebellum-associated (**Fig. 1**). For example, the locomotor group combined the nMLF mask from mapzebrain (Kunst et al., 2019; Shainer et al., 2023) with Mauthner-system masks from Z-Brain (Randlett et al., 2015). We added the mesencephalic locomotor region (Carbo-Tano et al., 2023) and prepontine long-latency C-start population (Marquart et al., 2019) from published localizations. For other reticulospinal populations, the activity patterns described by (Lau et al., 2025) guided masks for forward swimming, strong turns, and orienting towards prey. The forward-swim mask combined Z-Brain reticulospinal parcels, whereas the strong-turn mask was obtained by thresholding mapzebrain *vsx2* reporter expression, informed by studies of V2a reticulospinal neurons (Collins et al., 2025; Kimura et al., 2013). The prey-orienting mask followed published neuronal localizations (Gahtan et al., 2005). **Supplementary Table 1** provides the sources and construction details for all masks.

To place the zebrafish atlas in a vertebrate context, we illustrated the proposed mammalian counterparts schematically using mouse anatomy (**Fig. S1**), with supporting evidence and caveats detailed in **Supplementary Table 1**. Assignments integrated functional role, rhombomeric position, molecular identity, connectivity, and developmental lineage, where available. Strong correspondences include the oculomotor and trochlear nuclei (nIII and nIV) (Greaney et al., 2017; Higashijima et al., 2000; Wullimann et al., 1996), the abducens nucleus (nVI) (Brysch et al., 2019; Dowell et al., 2025), the vestibular nuclei (Liu et al., 2022; Straka and Baker, 2013), the locus coeruleus (Guo et al., 1999; Morin et al., 1997), the cerebellar cortical system (Bae et al., 2009; Hashimoto and Hibi, 2012), and the inferior olive (Apps and Garwicz, 2005; Chen et al., 2018; Félix et al., 2024; Yang et al., 2022). Not all functional regions have a direct mammalian homologue; the medial octavolateralis nucleus (MON), for example, belongs to an anamniote-specific sensory system (Bell, 2002; Valera et al., 2021; Vanwalleghem et al., 2020). The respiratory rhythm remains an open correspondence: no cell-resolved counterpart of the mammalian preBötzinger complex or RTN/parafacial system has yet been identified in zebrafish larvae (Del Negro et al., 2018; McArthur et al., 2023; Pan et al., 2019). Together, these comparisons place the zebrafish hindbrain atlas within a vertebrate framework while distinguishing well-supported correspondences from regions whose mammalian counterparts remain uncertain or absent (**Fig. S1**, **Supplementary Table 1**).

Having assembled this reference, we next asked whether unsupervised analysis of whole-hindbrain co-activity could recover established functional populations and reveal additional reproducible ensembles, without using anatomical or functional labels to define them.

### Unsupervised extraction of conserved synchronous ensembles

To examine hindbrain functional organization across the whole structure simultaneously, we recorded calcium activity with light-field microscopy (LFM), which captures the entire hindbrain in a single camera exposure (Aimon et al., 2019; Broxton et al., 2013). We imaged a pan-neuronal, nuclear-localized jGCaMP8s line (*elavl3:H2B-jGCaMP8s*) at 5 Hz. We also generated a complementary cytosolic line (*elavl3:jGCaMP8s*) for higher-rate imaging, acquiring recordings at 5–20 Hz. During neural imaging, we simultaneously tracked tail movements across the dataset, together with eye, pectoral-fin, and mouth movements in recordings where these were visible (**Fig. 2A**; **Methods**). We primarily recorded spontaneous behavior, either in darkness or in a closed-loop virtual-reality environment in which swimming updated a naturalistic visual scene (**Video S1**). We also presented moving gratings to elicit the optomotor response (**Fig. 2A**).

**Figure 2.**
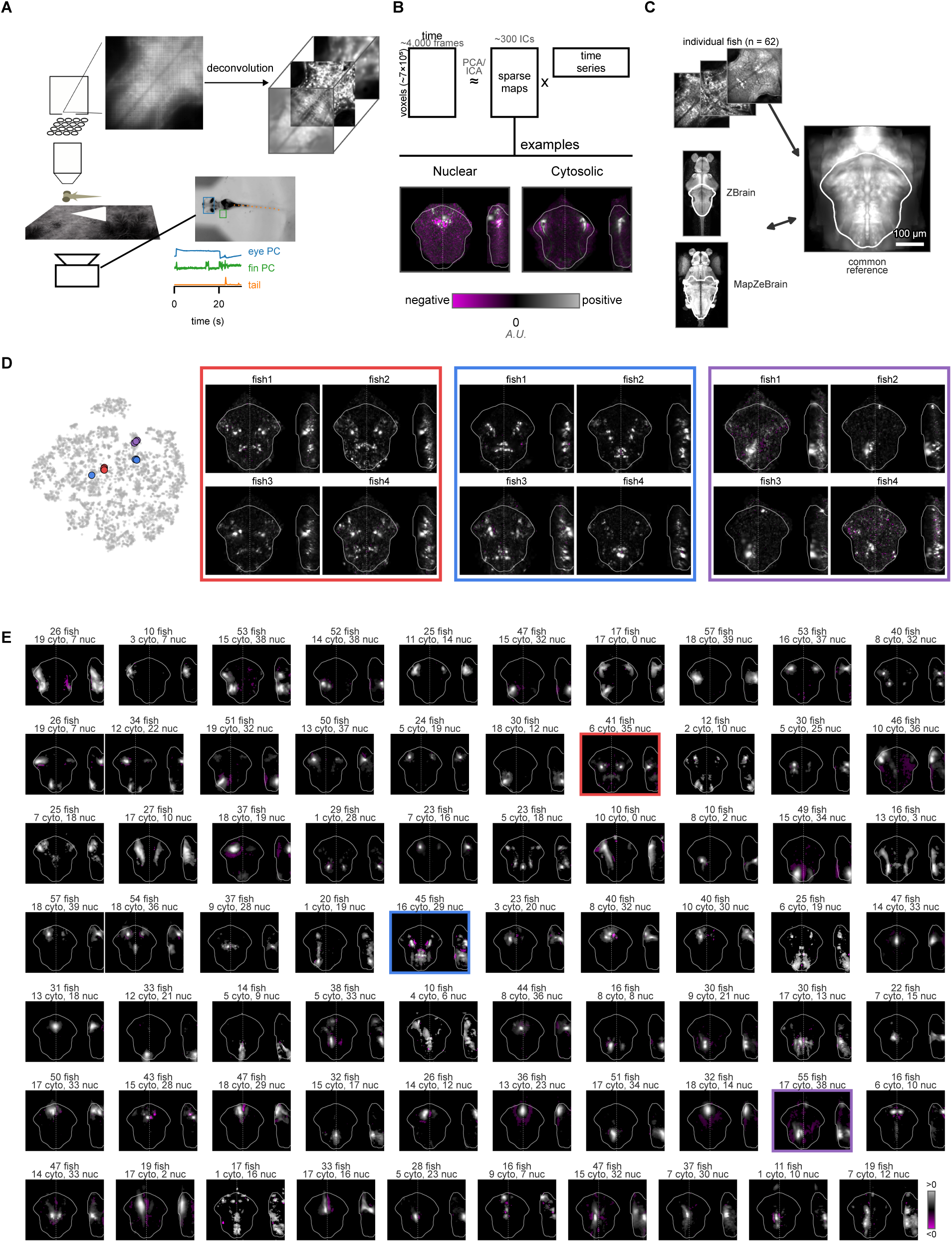
Unsupervised extraction of reproducible synchronous ensembles from whole-hindbrain activity. **A,** Whole-hindbrain imaging and behavioral tracking. Larvae were head-fixed with the eyes, pectoral fins, tail, and mouth free to move. During closed-loop trials, we updated the visual scene according to swimming movements estimated from tail tracking, providing visual feedback (**Methods**). Light-field microscopy captured the entire hindbrain in each frame, and volumes were reconstructed by deconvolution at a voxel size of 3 x 3 x 3 µm^3^ (Aimon et al., 2019; Broxton et al., 2013). Representative traces show PCA-derived eye and pectoral-fin signals and tracked tail-tip deflection, recorded simultaneously with neural activity (**Methods**). **B,** Unsupervised decomposition of hindbrain activity. For each recording, PCA/ICA decomposed the voxel-by-time matrix (∼7 × 10^5^ voxels and ∼4,000 frames) into ∼2,500 spatial maps and their associated time series; filtering retained approximately 300 ensembles per recording. Example ensembles from the nuclear-localized and cytosolic jGCaMP8s lines are shown. Within each example, gray and magenta show the positive and negative parts of the same ensemble’s spatial map. The cytosolic example extends into an associated fiber tract and neuropil. **C,** Registration to a common reference space. We registered individual fish anatomies with ANTs to a reference constructed by averaging anatomical volumes from several light-field recordings (**Methods**). We also aligned the Z-Brain and mapzebrain atlases to this reference. **D,** Visualization of ensemble similarity across fish. Each point in the t-SNE plot represents one ensemble extracted from an individual recording; the embedding visualizes similarity among ensembles using the spatial features employed for cross-fish clustering. Each colored frame shows spatial maps of matched ensembles from four different fish, in dorsal and lateral projections, with corresponding points highlighted in the same color. The bottom-right ensemble in the blue frame has a spatial map similar to the other three but lies apart from them in the embedding. This example illustrates a case in which manual curation was necessary. **E,** Atlas of reproducible hindbrain ensembles. Cross-fish average spatial maps for the 70 curated clusters with at least 10 fish (gray, positive weights; magenta, negative; hindbrain outline and midline overlaid). All ensembles were distinguishable in leave-one-fish-out cross-validation except two pairs with at least 15% reciprocal misclassification, shown without an intervening gap. Annotations above each map give the numbers of contributing fish from the cytosolic and nuclear-localized lines, respectively.

We used an unsupervised decomposition—principal component analysis (PCA) followed by spatial independent component analysis (ICA) (Aimon et al., 2023, 2019; Beckmann and Smith, 2004)—to extract sparse co-active ensembles from the neural activity data (**Fig. 2B**; **Methods**). Each ensemble thus comprises a spatial map of coordinated activity and an associated time series. Unlike methods that explicitly model spatially compact sources to segment somata or neurites (e.g., constrained non-negative matrix factorization, CNMF (Giovannucci et al., 2019)), this decomposition imposes no spatial-locality constraint, yet recovered ensembles were predominantly spatially localized. In the cytosolic line, ensembles additionally resolved their associated tracts and neuropil for some clusters (**Fig. 2B**).

To test whether these ensembles reflect shared hindbrain organization rather than per-fish idiosyncrasy, we registered all fish into a common reference brain and also aligned them to the Z-Brain and mapzebrain atlases (**Fig. 2C**) (Avants et al., 2008; Kunst et al., 2019; Randlett et al., 2015). We then clustered ensembles across fish using intensity profiles of the spatial weights, before refining clustering by manual curation (**Methods**). For visualization, we embedded the ensembles extracted from individual recordings using t-SNE (Van der Maaten and Hinton, 2008) (**Fig. 2D**). Ensemble maps were highly reproducible across individuals: clustering grouped ensembles from many different fish into clusters with closely matching maps (**Fig. 2D, E**). Across the cohort, we recovered 70 reproducible clusters, each represented in at least 10 distinct fish (medians: 72 ensembles and 35 fish per cluster; **Fig. 2E**, **Table 1**). Most ensembles were detected in both the cytosolic and nuclear-localized lines, while a subset appeared predominantly or exclusively in one (**Fig. 2E**, **Fig. S2**). We next assessed whether spatial features could predict the curated cluster assignments in held-out fish. Using leave-one-fish-out cross-validation, a classifier trained on the remaining fish assigned individual ensembles to one of the 70 curated clusters with 54% balanced accuracy (95% CI 51–57%; chance 1.4%; **Fig. 2E**). For each held-out fish, the reference map for each cluster was recalculated using only ensembles from the remaining fish (**Fig. 2E**). Together, these results show that the extracted ensembles recur across fish and can be recognized in animals not used to define them.

We next asked whether these ensembles could also be recovered from independently acquired data using different approaches. We re-analyzed four publicly available whole-brain light-sheet datasets (Chen et al., 2018; Lueckmann et al., 2025; Marquez-Legorreta et al., 2026; Van Der Plas et al., 2023), using either cell-level or voxel-level representations and distinct methods to identify functional groups. For the Lueckmann et al. and Marquez-Legorreta et al. datasets, we applied PCA followed by spatial ICA to voxel-level and cell-level data, respectively; for the Van Der Plas et al. and Chen et al. datasets, we used the authors’ released functional groups, obtained with compositional restricted Boltzmann machines and k-means correlation clustering, respectively. We first registered all ensembles obtained from these datasets to our reference space. We then matched them to our cluster averages using a common spatial-similarity measure followed by manual curation (**Fig. S3**, **Methods**). Of the 59 ensembles eligible for cross-dataset comparison—the 70 clusters displayed in **Fig. 2E** minus 11 whose members are predominantly (≥80%) from the cytosolic jGCaMP8s line and therefore not recoverable by the methods using only nuclear GCaMP—51 (86%) were recovered in at least one of the four datasets, 41 in at least two, 27 in at least three, and 14 in all four (**Fig. S3**). Ensembles recovered across methods are unlikely to arise from assumptions specific to any single decomposition.

Together, these analyses reveal a reproducible repertoire of hindbrain ensembles, including a core set recovered across fish, reporter lines, imaging modalities, and analysis methods. The resulting atlas provides spatially defined functional populations that can be compared across studies. These populations were identified from co-activity, leaving open how they relate to anatomical and molecular organization. We next addressed this question by comparing their spatial maps with anatomical regions and gene-expression patterns.

### Unsupervised extraction of expression clusters overlap with functional ensembles

Having identified reproducible functional ensembles from co-activity, we next asked whether their spatial organization corresponded to patterns of gene expression. We applied the same unsupervised decomposition used on the activity data to 747 registered reference expression maps from the mapzebrain atlas. These maps included transgenic and enhancer-trap reporter lines, HCR/ISH gene probes, and antibody stains. We excluded spatially diffuse components whose high-amplitude spatial footprint covered a large fraction of the hindbrain (**Methods**), leaving 658 sparse expression components for comparison with the functional ensembles (**Fig. 3A**). Each expression component delineated populations sharing a characteristic expression signature, but not necessarily their full expression profile; these shared signatures therefore do not define molecular cell types.

**Figure 3.**
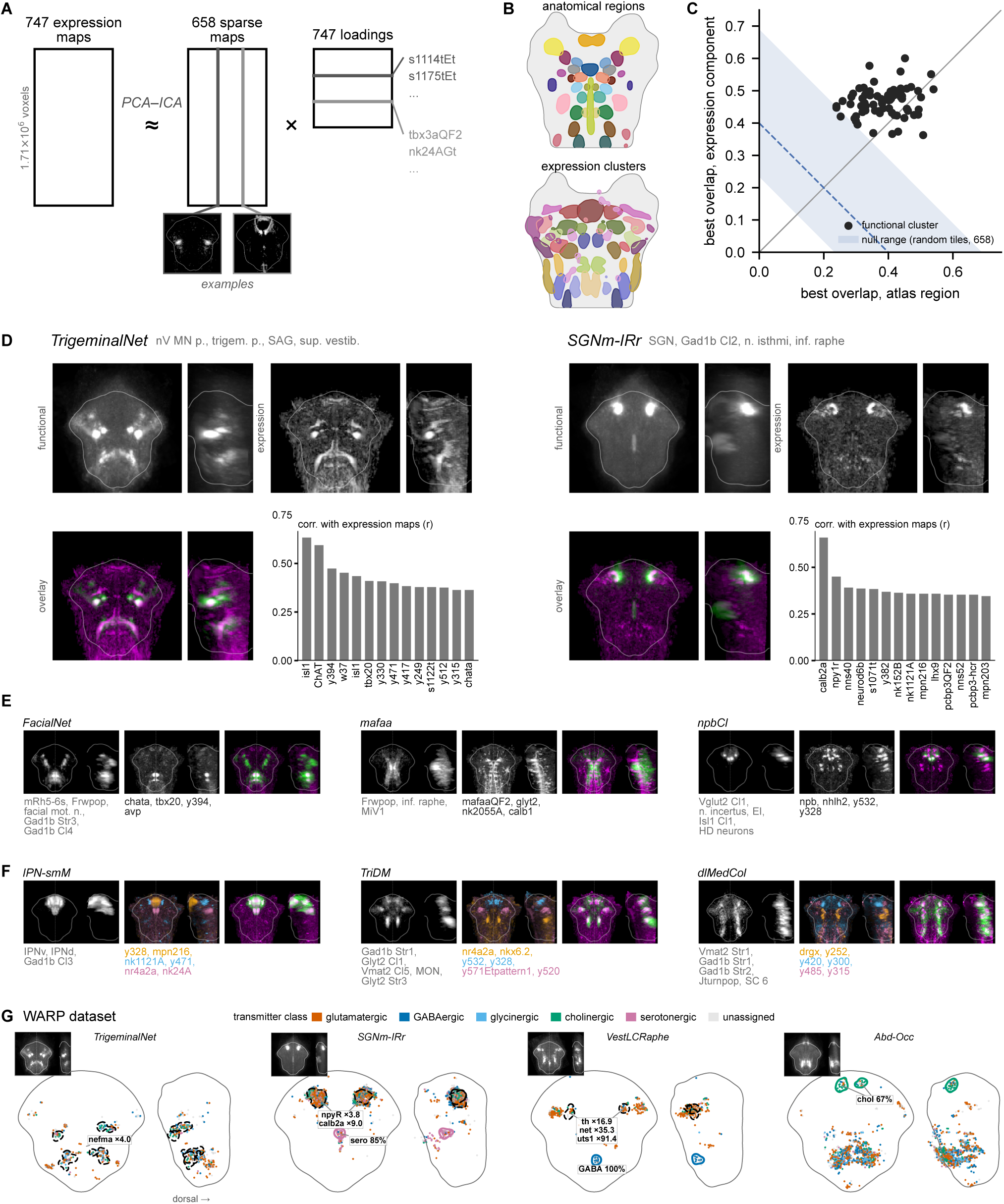
Expression-derived ensembles show strong spatial correspondence with functional ensembles. **A,** PCA–ICA decomposition of 747 registered reference expression maps into 746 components. We retained 658 sparse spatial components after the exclusion of 88 diffuse or noise-like components. Example components and their loadings onto the source maps are shown below. **B,** Whole-depth projection of 20 representative atlas regions (top) and 20 retained expression components (bottom, thresholded, see **Methods**) in the common light-field reference space; colors distinguish individual regions or components. **C,** Best spatial overlap (cosine similarity) of each functional cluster with an atlas region (x) versus with a retained expression component (y). Expression provides the stronger match for 58 of 70 functional clusters. The shaded blue band spans the best overlaps obtained with 658 arbitrary bilateral tiles of the hindbrain; the dashed line marks their mean. **D,** Expression matches for *TrigeminalNet* and *SGNm-IRr* functional clusters. The functional map, best-matching expression component, and their overlay are shown in dorsal and side views; gray text next to each name lists up to four atlas regions of the ensemble, ranked by overlap (**Methods**), and bar plots rank the 14 source expression maps most correlated with the matched expression component. **E,** Additional functional-to-expression matches. For each example, the functional map, matched expression component, and overlay are shown in dorsal and side views; gray labels identify overlapping atlas regions and black labels the four source maps most correlated with the component (**Methods**). **F,** Examples in which functional ensembles are captured by multiple expression components, selected by greedy residual matching (**Methods**). Each component is shown in its own color and overlaid with the functional map; the colored labels list the two source maps most correlated with each component. In the D to F overlays, functional maps are green and expression components are magenta. Region abbreviations: Cl, cluster; Str, stripe; MN, motor neurons; mot. n., motor nucleus; n., nucleus; trigem. p., posterior trigeminal motor nucleus; sup. vestib., superior vestibular nucleus; inf. raphe, inferior raphe; SC, spinal cord; EI, evidence integrator. All maps are displayed in the common reference space. The functional maps in D show the unthresholded across-fish average; in E, F and all overlays, functional maps are thresholded at 2.5 standard deviations of their non-zero voxels, and expression components are background-subtracted (**Methods**). **G**, Transmitter identity and gene expression of four clusters at cellular resolution in the WARP dataset. The high-loading neurons of every WARP ensemble assigned to the cluster, pooled across up to six larvae and restricted to the functional hindbrain mask, are colored by transmitter class measured by multiplexed HCR in the same cells; unassigned neurons are light gray. Left, dorsal view (rostral up); right, side view (dorsal to the right); the gray line outlines the hindbrain within the functional mask, and the inset shows the light-field cluster average. Colored outlines mark the largest glycinergic or cholinergic subcluster, the GABAergic subcluster of *VestLCRaphe* and the serotonergic subcluster of *SGNm-IRr*, each labeled with its share of typed cells. Dashed black outlines mark clusters of neurons positive for *nefma*, *npyR* or *th*, labeled with the fold enrichment over the hindbrain mean (for *VestLCRaphe* also *net* and *uts1*, for *SGNm-IRr* also *calb2a*, over the same clusters) (**Methods**).

To compare functional ensembles with anatomical and molecular reference maps in the same coordinate system, we transformed masks from the mapzebrain and Z-Brain atlases (along with additional masks from **Fig. 1**) into our reference space and visualized them alongside the expression-derived components (**Fig. 3B**). We also used these correspondences to assign descriptive names to the functional ensembles (**Table 1**). When an ensemble closely matched a known anatomical population, we named it for that location or circuit, as for *HBO* (a.k.a. ARTR). Other names summarize the ensemble’s spatial distribution across multiple regions, as for *Abd-Occ*, *VestLCRaphe* and *SGNm-IRr*. When a molecular pattern provided the clearest correspondence, we used the associated marker or transgenic label, as for *isl1*. The names are descriptive and do not imply exact anatomical or molecular identity.

We next asked whether expression-derived components or atlas regions more closely matched the spatial organization of the functional ensembles. We compared each functional ensemble with the expression-derived components and with 178 atlas regions from the mapzebrain and Z-Brain atlases, and **Fig. 1**, including molecularly defined masks (**Methods**). Most expression components combined contributions from several reporter lines and HCR maps, indicating composite molecular signatures (**Fig. 3A,D**). Expression-derived components provided a stronger spatial match than the atlas regions for 58 of 70 clusters (**Fig. 3C**) and exceeded the best overlap with arbitrary bilateral tiles of the hindbrain for all 70 clusters (**Fig. 3C**).

Several functional ensembles closely matched individual expression-derived components in their spatial organization. *TrigeminalNet* matched a component with a cholinergic-motor signature spanning posterior trigeminal and facial motor regions, whereas *SGNm-IRr* matched a component linking the medial part of the secondary gustatory nucleus with an inferior-raphe domain (**Fig. 3D**). Additional one-to-one matches occurred throughout the hindbrain (**Fig. 3E**). Other ensembles, including *IPN-smM, TriDM*, and *dlMedCol*, were better captured by combinations of several expression-derived components (**Fig. 3F**), suggesting that coherent functional activity can span multiple molecularly distinguishable populations. All functional clusters and their matches are listed in **Table 1**.

To examine the molecular composition of these ensembles at cellular resolution, we used the WARP dataset (Marquez-Legorreta et al., 2026), in which neural activity and multiplexed HCR measurements were obtained from the same neurons. For each ensemble, we pooled the neurons with the strongest ensemble loadings from all matched WARP ensembles across up to six larvae (**Fig. 3G**, **Fig. S4**, and **Methods**). The pooled neurons often comprised several transmitter classes, and within them we identified spatially localized subclusters enriched for individual transmitter classes or genes (**Fig. 3G**). *TrigeminalNet* neurons were enriched for the cholinergic marker *chata* (7.9-fold over the hindbrain mean) and *nefma*-positive neurons formed local clusters (4.0-fold), consistent with motor neurons. *SGNm-IRr* contained a spatially isolated subcluster in the inferior raphe that was 85% serotonergic and anterior clusters of *npyR*-positive neurons (3.8-fold) enriched for *calb2a* (9.0-fold). *VestLCRaphe* contained a compact medial subcluster in the inferior raphe that was entirely GABAergic, and its anterior *th*-positive clusters (16.9-fold) co-expressed *net* (35-fold) and *uts1* (91-fold), marking them as the noradrenergic locus coeruleus. The anterior cholinergic cluster of *Abd-Occ* (67% cholinergic) lay at the position of the oculomotor or trochlear nuclei. Because transmitter identity and gene expression were not used to define or match the ensembles, they provide independent molecular support for the spatial assignments. See **Table 1** for all cluster compositions, and **Fig. S4** for additional maps.

Together, these results show that distinctive spatial expression patterns correspond closely to independently identified co-active ensembles, linking coordinated activity to shared aspects of molecular identity.

### Functional ensembles and hindbrain gene expression share strong bilateral symmetry

The close spatial correspondence between functional and expression-derived components led us to ask whether they also shared the hindbrain’s prominent left–right organization (Collins et al., 2025; Hildebrand et al., 2017; Metcalfe et al., 1986). We quantified the self-symmetry of each component by comparing its spatial map with its left–right reflection (**Methods**). In the functional decomposition, bilateral organization could take four forms: (1) a self-symmetric ensemble spanning both sides with the same sign; (2) a self-antisymmetric ensemble spanning both sides with opposite signs; (3) separate left- and right-sided ensembles forming a co-active mirrored pair; or (4) separate left- and right-sided ensembles forming an anti-active mirrored pair (**Fig. 4A**). For lateralized ensembles, we searched within the same recording for a spatially mirrored partner and used the correlation between their time series to distinguish co-active from anti-active pairs. Ensembles concentrated around the midline were classified as midline, whereas those lacking any of these forms of bilateral correspondence were classified as asymmetric.

**Figure 4.**
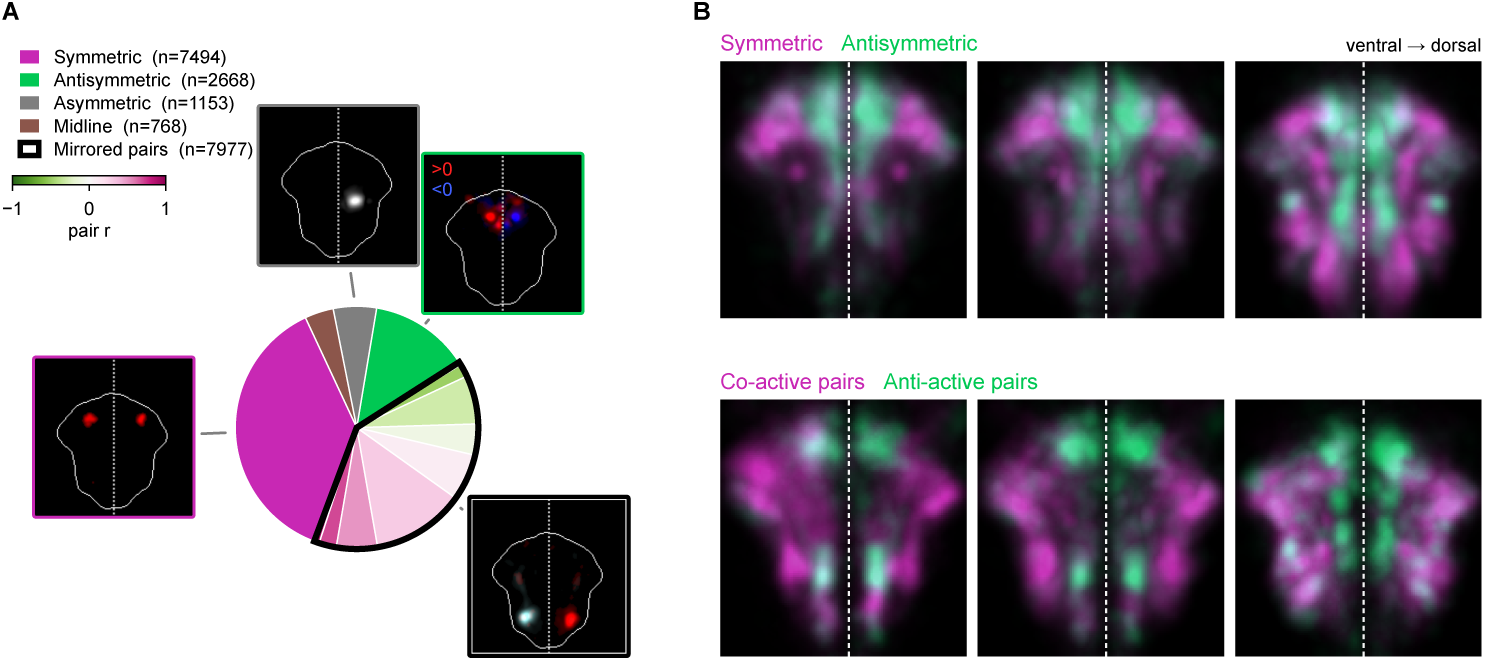
Hindbrain functional ensembles are bilaterally organized. **A,** Distribution of individual functional ensembles (n = 20,060, pooled across 68 recordings from 50 larvae) across left–right symmetry categories. For symmetry measurements, maps were spatially binned from 3-µm to 24-µm voxels (**Methods**). Ensembles were classified as bilaterally symmetric (magenta; mirrored spatial distributions with the same sign on both sides), bilaterally antisymmetric (green; mirrored distributions with opposite signs), mirrored pairs (black-outlined wedge; lateralized ensembles whose left–right reflection matched another ensemble from the same recording), asymmetric (gray; lateralized ensembles without a mirrored partner), or midline (brown). For mirrored pairs, shading indicates the Pearson correlation between the two ensemble time series, ranging from anti-active pairs with negative correlation to co-active pairs with positive correlation. A representative ensemble is shown for each category except midline. Symmetric and antisymmetric examples show positive values in red and negative values in blue (key in the antisymmetric example). The mirrored-pair example shows one ensemble in red and its partner in the color of their time-series correlation. **B,** Category-average maps resolved by depth, overlaid in pairs: symmetric (magenta) with antisymmetric (green) ensembles (top), and co-active (magenta) with anti-active (green) mirrored pairs with |r| > 0.3 (bottom). Columns show three successive depth ranges from ventral (left) to dorsal (right); each is the maximum projection of the category mean over that range.

Hindbrain gene-expression components were strongly self-symmetric (96% of 658 non-noise components; 2% antisymmetric, 2% midline, 0% asymmetric). Across 20,060 functional ensembles, 37% were self-symmetric, 13% self-antisymmetric, 6% asymmetric, 4% midline, and 40% belonged to mirrored pairs (27% co-active, 13% anti-active) (**Fig. 4A**). These categories also differed in their mediolateral distribution: self-antisymmetric ensembles and anti-active mirrored pairs lay closest to the midline (center-of-mass 49 and 54 µm off-midline), whereas self-symmetric ensembles and co-active mirrored pairs extended further laterally (63 and 79 µm) (**Fig. 4B**). Together, these results show that the strong bilateral organization of hindbrain gene expression is paralleled in activity.

### Reproducible hindbrain ensembles track distinct sensorimotor variables and temporal features

Having established the spatial, molecular, and bilateral organization of the ensembles, we next examined their relationships to sensory input and behavior. We compared each ensemble’s activity with visual stimulation and movements of the eyes, tail, pectoral fins and mouth (**Methods**).

Two clusters, *aHb* and *npbCl*, showed direction-dependent responses to whole-field visual motion (**Fig. 5A**; fish-level t-test, FDR q < 0.05). The *aHb* response was consistent with previously described visual-motion responses in the anterior hindbrain (Bahl and Engert, 2020; Dragomir et al., 2020). Although modulated by visual stimulation, these clusters could also be extracted from recordings in darkness, indicating that patterned visual input was not required for their coordinated activity. Across the atlas, no cluster differed significantly in detection frequency between recordings in darkness and those with visual stimulation (**Fig. S5A**).

**Figure 5.**
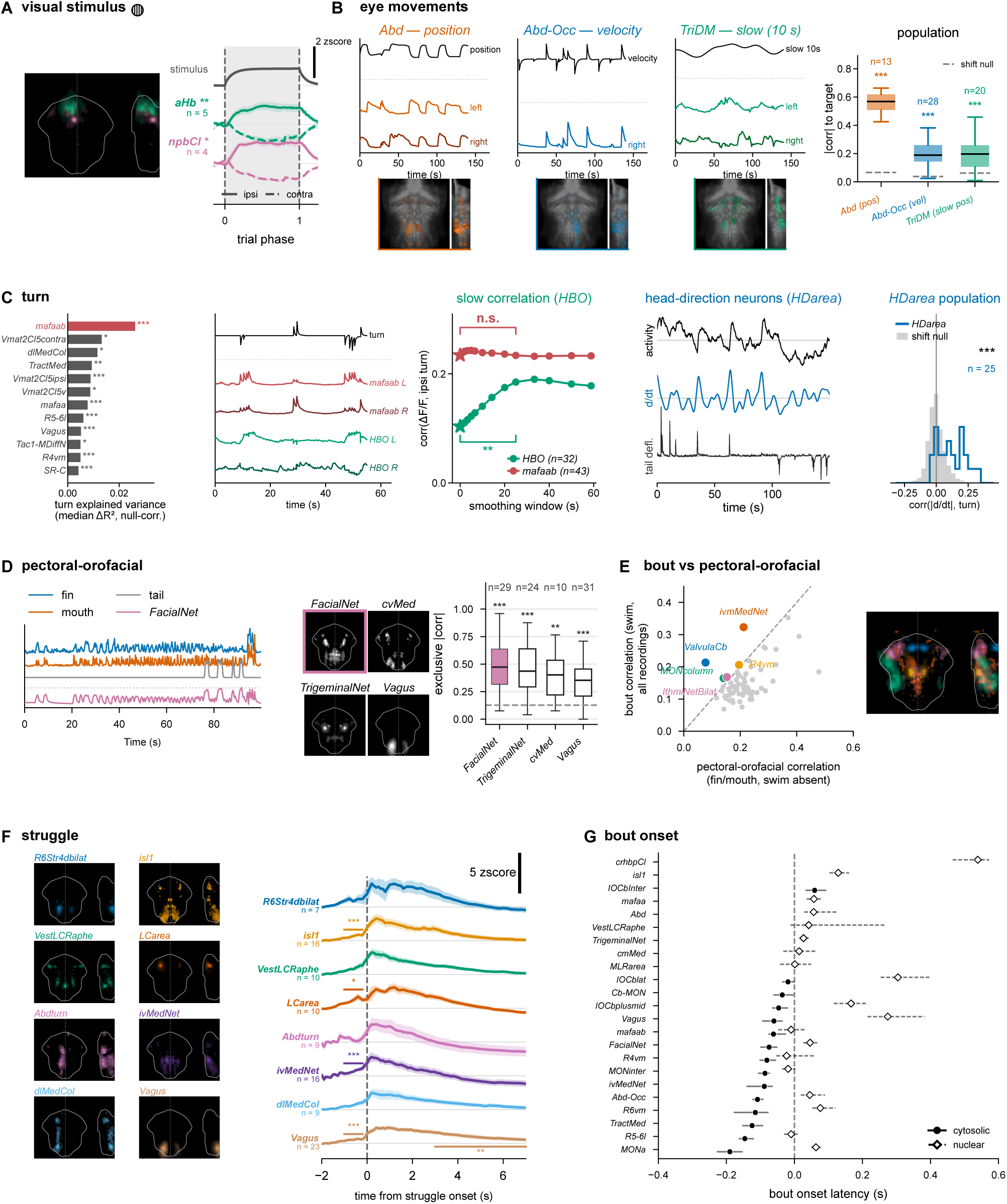
Hindbrain ensembles differ in their sensorimotor associations and response dynamics. **A,** Visual stimulus (left- and right-moving gratings). Left: footprints of the two clusters showing a significant difference between ipsilateral and contralateral whole-field motion. Right: ensemble activity during whole-field motion, time-normalized from motion onset (0) to offset (1), split into motion toward the ipsilateral (solid) and contralateral (dashed) side of each ensemble’s hemisphere (mean ± s.e.m. across fish; baseline-subtracted z-scored ΔF/F; nuclear reporter line). . Gray line, stimulus regressor obtained by convolution with the jGCaMP8s response kernel. Significant ipsi–contra differences were identified by fish-level t-tests over the first 80% of the motion window. **B,** Eye-movement-related signals are decomposed into position, velocity, and slow signals. Three ensembles capture distinct eye-movement codes: *Abd* tracks eye position, *Abd-Occ* tracks eye velocity (d/dt of position), and *TriDM* tracks a smoothed eye position signal (∼10 s low-pass of position). For each, the left and right example ensembles (colored) are shown over the matched eye regressor (gray); all three examples are from the same recording. Cluster footprints are shown below (top + lateral views). Right: absolute correlation of each cluster’s ensembles with its target eye signal (*Abd*, eye position; *Abd-Occ*, eye velocity; *TriDM*, smoothed eye position), averaged within each fish (n = fish); dashed lines, circular-shift chance level. **C,** Turn-related ensembles show distinct temporal relationships with turning. Left: turn coupling for the clusters with a significant effect, ranked by the median across fish of the null-corrected turn ΔR² (fish-level one-sided Wilcoxon, FDR q < 0.05); the fast turn-execution cluster *mafaab* is highlighted in red. Middle: turn coupling as a function of smoothing window, mean across fish (n = fish); brackets compare the native signal with the ∼25 s window (FDR-corrected); *HBO* (ARTR) correlation increases with smoothing, whereas *mafaab* is already maximal at short windows. Right two panels: *HDarea*’s activity derivative tracks turning (consistent with a heading-direction representation). Component activity, its derivative, and tail deflection are shown; the distribution of per-fish mean corr(|d/dt|, turn) against a circular-shift null (n = fish). **D,** Pectoral fin and mouth (orofacial) movements. Example *FacialNet* ensemble shown against simultaneous fin, mouth, and tail movements; cluster footprints (4 most correlated clusters); boxplot ranks clusters by orofacial exclusive |corr| (the larger of the fin- and mouth-specific correlations), one value per fish (n = fish); dashed line, chance; asterisks, fish-level one-sided Wilcoxon against chance, FDR-corrected across all tested clusters. **E,** Bout-specific (non-orofacial) ensembles. Bout versus orofacial-only correlation (fin/mouth exclusive |corr|). Clusters above the identity line, whose bout coupling exceeds their orofacial coupling, are highlighted in color: *ivmMedNet*, *ValvulaCb*, *R4vm*, *MONcolumn* and *IthmiNetBilat*. Right: overlaid footprints of the highlighted clusters, each in its scatter color. **F,** Peri-struggle-onset activity (mean ± s.e.m. across fish; time 0 = onset of a manually verified struggle) for the seven clusters with the largest response 0–1.5 s after onset, and *Vagus*. Left: footprint of each cluster in its trace color. Brackets mark significant comparisons: −1 to −0.2 s, a larger rise before struggles than before normal swim bouts; 3–7 s (Vagus), activity above baseline (fish-level Wilcoxon, Holm-corrected across the 16 tests; unmarked comparisons were not significant). **G,** Bout-onset latency (time to half-maximum of the swim-onset-aligned response; 68% bootstrap interval over ensembles) for clusters with a significant bout-onset response, ranked earliest to latest, for the cytosolic (filled circles, solid) and nuclear (open diamonds, dashed) GCaMP lines. Statistics: *p < 0.05, **p < 0.01, ***p < 0.001 after the correction stated per panel. Unless stated otherwise, ensembles were averaged within each fish and tests were performed across fish; n is the number of fish and error bands are s.e.m. across fish; in G, the unit is the ensemble.

Eye-related ensembles followed different kinematic signals. *Abd* tracked conjugate eye position, *Abd-Occ* followed eye velocity, and *TriDM* tracked a slowly varying eye-position signal obtained by smoothing over approximately 10 s (**Fig. 5B**). These different response patterns were apparent in ensembles recorded simultaneously in the same animal. For each of the three clusters, correlations with the corresponding eye signal exceeded a circular-shift null across fish (FDR q < 0.001).

Turning was also associated with distinct activity dynamics. We first ranked turn coupling by the decrease in explained variance after removing the signed turn regressor from a joint model containing the available stimulus and behavioral regressors, with correction against a circular-shift null (grouped ΔR²; **Methods**). By this measure, *mafaab* showed the strongest turn coupling (**Fig. 5C**). Its activity followed discrete turning events, whereas *HBO* followed slower variations in turning. Smoothing the turn signal over tens of seconds increased its correlation with *HBO* activity but produced little change in *mafaab* coupling. *HDarea* showed a different relationship: the magnitude of its activity derivative correlated with turning above the circular-shift null, associating turns with changes in ensemble activity (**Fig. 5C**). This is consistent with a representation of heading (Petrucco et al., 2023).

Because movements of different body parts frequently occurred together, swimming correlations alone could not resolve their individual associations with ensemble activity. Behavioral cross-correlograms showed particularly strong coordination between fin and mouth movements, which were also closely coupled to tail movements (**Fig. S5B**). Because fin and mouth were strongly correlated with each other, we did not attempt to assign ensembles separately to the two effectors. Nevertheless, *FacialNet* activity followed repeated fin and mouth movements during a long interval without tail movement (**Fig. 5D**). To quantify these associations, we measured absolute correlations during periods when the corresponding effector moved while the tail was immobile and summarized each ensemble by the larger of its available fin- and mouth-specific correlations (**Methods**). *FacialNet*, *TrigeminalNet*, *cvMed* and *Vagus* showed the strongest coupling by this measure, identifying ensembles associated with fin or mouth movements in the absence of swimming (**Fig. 5D**; each above chance, FDR q < 0.01).

The converse comparison identified ensembles with stronger swim associations. Comparing swim correlations across all recordings with isolated fin/mouth correlations highlighted *ivmMedNet* (a cluster to the medial anterior part of the facial nucleus) and *ValvulaCb* as more strongly associated with swim bouts (**Fig. 5E**). *R4vm*, *MONcolumn* and *IthmiNetBilat* showed the same tendency. Thus, tracking additional body parts distinguished ensembles associated with isolated fin or mouth movements from ensembles more strongly associated with swimming.

Struggle-related ensembles differed in their timing and persistence. Activity in *isl1*, *ivMedNet*, *LCarea* and *Vagus* rose more in the second before struggle onset than before normal swim bouts (**Fig. 5F**; fish-level Wilcoxon, Holm-corrected). *Vagus* was the only cluster whose activity stayed elevated 3–7 s after onset, consistent with the response of noradrenergic neurons in this region (Mu et al., 2019).

Finally, we compared bout-onset response timing across the ensemble atlas. Responses included both increases and decreases in activity, with turn-direction-dependent responses in ensembles such as *mafaab* and *Abd-Occ* (**Fig. S5C**). By matching ensembles across reporter lines and aligning their activity to bout onset, we could also compare calcium-response dynamics and provide an empirical benchmark for the slower kinetics of nuclear-localized H2B-jGCaMP8s relative to cytosolic jGCaMP8s. For ensembles with sufficiently large onset-aligned responses, we measured the first time at which the response reached half its maximum absolute magnitude, analyzing the two reporter lines separately (**Methods**). These latencies spanned a graded range, with some ensembles reaching half-maximum before detected swim onset and others afterwards (**Fig. 5G**).

Together, these analyses recover established sensorimotor populations and reveal new brain– behavior relationships, including ensemble activity associated with fin and mouth movements in the absence of swimming. Ensembles identified from co-activity alone thus differ in the sensory and motor variables they track, their coupling to fast and slowly varying behavioral signals, and their timing around movement onset.

### Activity-defined ensembles recover established populations and provide starting points for circuit discovery

We next asked how the activity-defined ensemble atlas relates to established hindbrain populations (**Fig. 1**) and what additional functional organization emerges from co-activity. Neither the PCA–ICA decomposition nor the cross-animal clustering used anatomical, molecular, or behavioral labels. To compare the resulting ensembles with the reference atlas, we organized them into the same eight functional and anatomical groups (**Fig. 6A**). Group assignments were based on measured sensorimotor associations (**Fig. 5**) or, when the relevant variables were not assessed, on spatial correspondence with established populations. For example, ensembles overlapping the medial octavolateralis nucleus were assigned to the mechanosensory group, although lateral-line responses were not directly tested.

**Figure 6.**
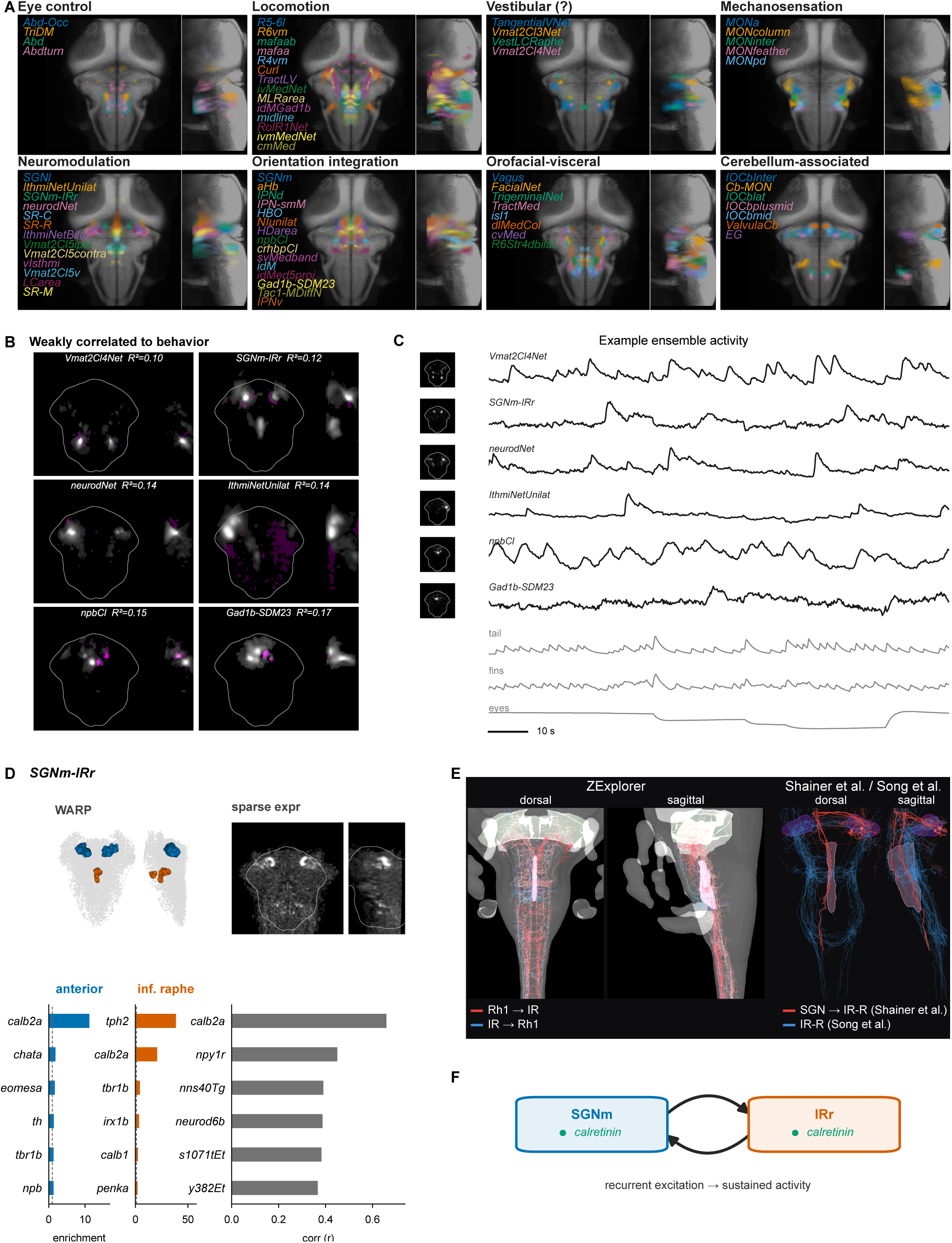
Activity-defined ensembles recover established hindbrain populations and provide functionally, molecularly, and anatomically constrained entry points for circuit discovery. **A,** Curated ensemble atlas organized into the eight functional and anatomical groups used in Fig. 1: eye control, locomotion, vestibular, mechanosensation, neuromodulation, orientation integration, orofacial–visceral, and cerebellum-associated. Ensembles were assigned according to their measured sensorimotor associations (Fig. 5) or, when the relevant variables were not assessed, their anatomical location. “Vestibular (?)” denotes ensembles located in the broader vestibular territory whose correspondence to individual vestibular nuclei and vestibular function remain unconfirmed. These assignments were made after ensemble extraction and cross-animal clustering. Maps show absolute ensemble averages across fish in mapzebrain space and shown as dorsal and lateral maximum-intensity projections, with each ensemble in a distinct color and labels color-matched to the maps. **B,** The six ensembles least explained by the measured stimulus and behavioral variables. For each ensemble, a lagged ridge model containing all available stimulus and behavioral regressors, with copies shifted from −3 to +3 s, was fitted to its activity and to its rate of change; each fit was scored as null-corrected R² (in-sample R² minus the mean R² after circular time shifts), and the ensemble took the larger value (**Methods**). Maps show cluster averages in dorsal and lateral projection (gray, positive weights; magenta, negative weights); labels give the cluster name and its median null-corrected R². **C,** Representative ensembles from the six weakly explained clusters in B, recorded simultaneously in one H2B-jGCaMP8s larva. Left, spatial map of the representative ensemble of the cluster in this recording; right, its z-scored activity. Gray traces show the corresponding jGCaMP8s-convolved swim, summed left- and right-pectoral-fin, and eye position regressors. Neural activity was structured but not visibly aligned with these measured variables. **D,** Molecular organization of *SGNm-IRr*. High-loading neurons pooled from WARP ensembles assigned to the cluster were grouped into an anterior-hindbrain group (blue) and a predominantly serotonergic inferior-raphe group (orange), each pooling spatial subclusters (**Methods**); gray shows all analyzed hindbrain cells. Bar plots show the six genes most enriched in each group relative to the hindbrain average. On the right, the sparse expression-derived component matched to *SGNm-IRr* in Fig. 3D (top) and the six source expression maps most strongly correlated with it (bars) are reproduced. *calb2a* was the most enriched gene in the anterior group, the second most enriched in the inferior-raphe group, and the source map most strongly correlated with the matched expression component. **E,** Anatomical evidence linking the anterior hindbrain to the right inferior raphe (each block shown in dorsal and sagittal views). Left, ZExplorer connectome query for reciprocal rhombomere-1–inferior-raphe projections (red, Rh1 → inferior raphe, mostly glutamatergic; blue, inferior raphe → Rh1). Rhombomere 1 and inferior-raphe masks are shaded. Right, single-neuron reconstructions from Shainer et al. (red, secondary gustatory nucleus neurons projecting to the right inferior raphe, SGN → IR-R) and *tph2*-positive inferior-raphe neurons from Song et al. (blue). SGN and inferior-raphe masks are shaded. **F,** Circuit model hypothesis summarizing **D** and **E**: *SGNm* (medial secondary gustatory nucleus) and *IRr* (right inferior raphe), each calretinin (*calb2a*)-positive, linked by reciprocal projections. These candidate reciprocal projections provide a potential substrate for coordinated activity between the two domains.

We recovered reproducible ensembles corresponding to a range of established populations. These correspondences were not always one-to-one: some named populations corresponded to several ensembles, while some ensembles spanned multiple named populations. Some of the latter formed distributed, multi-region ensembles spanning territories assigned to different functional groups in the literature-based atlas. *Abd-Occ* recapitulated the known coordination between abducens and oculomotor populations, whereas *TriDM*, *TrigeminalNet*, *VestLCRaphe* and *SGNm-IRr* linked spatially separated territories across different functional groups. These ensembles were highly reproducible across fish and were also recovered in independent datasets using different imaging and analysis approaches (**Fig. S3**). We also identified reproducible ensembles without a clear counterpart in the literature-based atlas.

To identify ensembles whose activity was poorly captured by the measured stimulus and behavioral variables, we fitted each ensemble’s activity and its rate of change with a lagged ridge-regression model containing all available stimulus and behavioral regressors, and scored each ensemble by the larger of the two null-corrected R² values (Methods). The six least explained clusters (0.10–0.17) occupied isthmic, nucleus incertus, dorsal-medulla and broader vestibular territories, and included a distributed ensemble spanning the anterior hindbrain and inferior raphe (**Fig. 6B**).

In a recording with a visual scene and simultaneous tail, pectoral fin, and eye tracking, ensembles from all six clusters showed prominent activity fluctuations that were not visibly aligned with the measured movements (**Fig. 6C**). Their dynamics ranged from sparse, high-amplitude transients in *neurodNet* and *IthmiNetUnilat* to slower fluctuations in *SGNm-IRr* and *npbCl*. All of these ensembles were also recovered in darkness (**Fig. S5A**), showing that their extraction did not require patterned visual stimulation.

We next asked whether molecular and anatomical reference data could help identify the populations underlying these weakly explained signals. We focused on *SGNm-IRr*, which spanned paired domains in the anterior hindbrain and a compact domain in the inferior raphe (**Fig. 6B,C**). In the cell-resolved WARP dataset, neurons contributing strongly to WARP ensembles matched to *SGNm-IRr* formed an anterior-hindbrain group and a predominantly serotonergic inferior-raphe group (**Fig. 6D**). *calb2a* was the most strongly enriched marker in the anterior group and, after *tph2*, in the inferior-raphe group. Independently, *calb2a* was the source expression map most strongly correlated with the expression-derived component matching *SGNm-IRr* (**Figs. 3D and 6D**). These two analyses therefore identified *calb2a* enrichment as a shared molecular feature of the co-active anterior-hindbrain and inferior-raphe populations.

We next asked whether anatomical projections could link the two co-active domains of *SGNm-IRr*. We queried ZExplorer (Du et al., 2026), a registered atlas of reconstructed single neurons, for projections between Rhombomere 1 and the inferior raphe. The rhombomere-1 neurons identified by this query had somata in the anterior territory occupied by *SGNm-IRr* and were mainly excitatory. Conversely, inferior-raphe neurons extended projections into rhombomere 1 (**Fig. 6E**, left). We then compared published reconstructions of secondary gustatory nucleus (SGN) neurons (Shainer et al., 2023) with those of serotonergic neurons in the rostral inferior raphe (IR-R; (Song et al., 2025)). This spatial comparison showed SGN neurites traversing the inferior-raphe territory and ascending IR-R projections reaching the anterior hindbrain (**Fig. 6E**, right). Together with the co-activity and shared *calb2a* enrichment of the two domains (**Fig. 6D**), these anatomical correspondences suggest a candidate bidirectional circuit (**Fig. 6F**), although synaptic connections between the participating neurons remain to be established.

## DISCUSSION

Here we establish an activity-based atlas of the larval zebrafish hindbrain, using the spatial organization of co-active ensembles to relate functional, molecular, anatomical, and behavioral measurements. We identify 70 clusters of ensembles that recur across animals, many of which are also recovered in independent datasets analyzed with different methods. Their spatial maps correspond to established functional populations and gene-expression patterns, reveal prominent bilateral and distributed organization, and distinguish populations associated with different sensory and behavioral variables. Ensembles weakly explained by the measured variables can nevertheless be investigated through their molecular and anatomical correspondences, as illustrated by *SGNm-IRr*. Together, these results establish reproducible ensemble maps as a common reference for relating complementary datasets and generating testable hypotheses about distributed hindbrain circuits.

Relating neuronal activity to predefined stimulus and behavioral variables provides an interpretable description of population function, but projects high-dimensional neural dynamics onto a limited set of external variables. In our recordings, even ensembles with strong direction-dependent visual responses were also recovered in darkness, indicating that patterned visual input modulated but did not define their coordinated activity. Activity-driven approaches provide a complementary view by identifying recurring population structure directly from neural activity, before assigning it a sensory or behavioral interpretation (Aimon et al., 2019; Mackevicius et al., 2019; Rubin et al., 2019; Van Der Plas et al., 2023). Here, defining ensembles from co-activity allowed us both to distinguish populations with broadly similar behavioral associations and to retain populations whose activity was weakly explained by the measured variables. Their spatial organization then provided a basis for comparison across animals and datasets and for identifying molecular and anatomical correspondences, even when their sensorimotor roles remained unresolved.

Many of the ensembles identified in our light-field recordings were also recovered in independent whole-brain light-sheet datasets ((Chen et al., 2018; Lueckmann et al., 2025; Marquez-Legorreta et al., 2026; Van Der Plas et al., 2023); **Fig. S3**). Corresponding spatial patterns emerged across distinct experimental and analytical approaches, supporting their use as a functional reference across datasets. This recovery is notable given substantial differences in experimental preparation, including pharmacological paralysis in three of the four comparison datasets, which may also contribute to incomplete recovery. Registered zebrafish atlases already provide common coordinates for comparing anatomy, gene expression, and neuronal morphology; reproducible functional ensembles add another level at which complementary datasets can be related. WARP (Marquez-Legorreta et al., 2026) links activity to molecular measurements in the same cells, whereas ZapBench (Lueckmann et al., 2025) links activity to electron microscopy of the same brain. Matching corresponding ensembles across these datasets therefore provides an ensemble-level bridge between molecular and circuit-level information that were not measured together.

Beyond the same-cell measurements in WARP, molecular correspondence was evident in the spatial organization of entire ensembles. Independent decomposition of registered transgenic and HCR expression maps yielded components that capture spatial patterns shared across multiple input maps rather than individual molecular cell types (Kunst et al., 2019; Shainer et al., 2023). These independently derived expression components showed close spatial correspondence with functional ensembles (**Fig. 3C**), linking the organization of co-activity to recurring structure in gene expression. This finding complements evidence that regional gene expression predicts interregional functional connectivity in zebrafish (Légaré et al., 2025), and that molecularly similar neurons exhibit stronger activity correlations in both zebrafish and mouse cortex (Bugeon et al., 2022; Marquez-Legorreta et al., 2026). Developmental organization offers one possible basis for this correspondence, as molecular identity and birth order are linked to neuronal position, morphology, connectivity, and recruitment during behavior in the zebrafish hindbrain (Kinkhabwala et al., 2011; Pujala and Koyama, 2019).

A further feature of the molecular correspondences was the prevalence of neuropeptide-related genes. Peptide ligands, precursors, or receptors were present among the molecular annotations of each of the six ensembles least explained by the measured stimulus and behavioral variables (**Fig. 6B**; **Table 1**), including *npb*/*tac1* in *npbCl*, *sst*/*cort* in *Gad1b-SDM23*, *cckb* in *IthmiNetUnilat*, and *npy1r* in *SGNm-IRr*. These signatures frequently occurred within spatial subpopulations that also had an assigned transmitter class, suggesting that peptide phenotype provides an additional molecular axis of ensemble organization. Neuropeptides are commonly co-transmitted with fast neurotransmitters and can signal outside classical synaptic specializations, often through G-protein-coupled receptors and on slower timescales (van den Pol, 2012). Peptide precursor and receptor expression also distinguish candidate sources from targets of modulation: for example, *npbCl* was independently matched to and enriched for *npb*, whereas *npy1r* expression in SGNm-IRr is consistent with previous identification of NPY-receptor-expressing neurons in the feeding-responsive SGN (Shainer et al., 2023). These observations raise the hypothesis that peptidergic modulation contributes to aspects of ensemble activity related to internal state that were not captured by our sensorimotor measurements.

The correspondence between functional and molecular maps also extended to their bilateral organization. Left–right pairing was prominent across these different arrangements, consistent with the bilateral organization of identified reticulospinal neurons and their axonal trajectories (Hildebrand et al., 2017; Metcalfe et al., 1986). This spatial symmetry nevertheless accommodated different relationships between the two sides. Whereas expression components were predominantly self-symmetric, functional ensembles included both self-symmetric and self-antisymmetric maps, together with separate mirrored pairs whose time series were co-active or anti-active (**Fig. 4A**). Opposing left– right activity is well established in the goldfish oculomotor integrator and the zebrafish HBO/ARTR, where it contributes to gaze control and directional biases in movement (Dunn et al., 2016; Wolf et al., 2017). Circuit studies show that interactions across the midline can support different forms of coordination: inhibitory competition contributes to left–right escape selection in zebrafish, whereas commissural projections are required for bilateral synchronization of the mouse inspiratory rhythm generator (Bouvier et al., 2010; Koyama et al., 2016). Our analysis extends this description beyond individual circuits and reveals a spatial bias: self-antisymmetric ensembles and anti-active mirrored pairs were concentrated closer to the midline than self-symmetric ensembles and co-active pairs (**Fig. 4B**). This distribution may reflect a more restricted spatial reach of inhibitory coupling than of the pathways supporting bilateral co-activation. In the cat visual cortex, inhibitory basket-cell connectivity is more local than excitatory connectivity (Stepanyants et al., 2009), while zebrafish spinal V1 interneurons inhibit motor and premotor targets locally despite extending axons over several segments (Sengupta et al., 2021). A similar restriction in the hindbrain could favor inhibitory competition between nearby populations on opposite sides of the midline, whereas more widely separated populations could remain co-active through longer-range excitation or shared inputs. Whether differences in the spatial reach of these pathways explain the observed medial bias could be tested by combining transmitter-annotated neuronal reconstructions with targeted measurements of functional connectivity.

Co-activity extended beyond the populations grouped within individual ensembles: distinct ensembles formed co-activity blocks across distant hindbrain areas (**Fig. S6**). Their shared activity nevertheless coexisted with differences in the sensory and behavioral signals they tracked (**Fig. 5**). Interpreting these associations required measuring multiple body parts, because eye, tail, fin, and mouth movements often occurred together; activity correlated with swimming could therefore also relate to accompanying movements (**Fig. S5B**). Even ensembles associated with the same effector differed in their temporal properties. Eye-related ensembles distinguished position, velocity and slowly varying signals, complementing the diversity of kinematic coding and persistence times described in hindbrain oculomotor circuits ((Brysch et al., 2019; Miri et al., 2011); **Fig. 5B**). Similarly, *mafaab* tracked discrete turns whereas *HBO* followed slower variations in turning, consistent with the established contribution of HBO/ARTR to directional biases across successive bouts ((Dunn et al., 2016); **Fig. 5C**). Response timing relative to swim onset provided another distinction among ensembles (**Fig. 5G**). These timing differences motivate tests of contributions to sensory integration and decision-making (Bahl and Engert, 2020; Dragomir et al., 2020), motor-command generation (Carbo-Tano et al., 2023), and processing of movement-related sensory feedback (Ahrens et al., 2012), functions already investigated in specific zebrafish circuits. Although calcium-response latencies alone cannot assign these roles, combining the ensemble maps with controlled sensory manipulations and targeted perturbations could distinguish populations involved in preparing, executing and monitoring movements. The atlas thus describes both which ensembles are co-active and how their individual relationships to behavior differ.

*TrigeminalNet* and *FacialNet* were coupled to fin or mouth movements in the absence of tail movement (**Fig. 5D**), suggesting possible roles in coordinating respiratory movements. This interpretation is consistent with coordinated opercular and pectoral-fin movements, together with rhythmic facial branchiomotor activity in larval zebrafish (McArthur et al., 2023; Zimmer et al., 2020). Recording their activity alongside resolved opercular, jaw and fin movements under controlled oxygen conditions could establish their relationship to respiration, while targeted perturbations could test whether individual populations coordinate these movements, generate their rhythm or convey motor commands. Identifying a rhythm-generating population would also provide a basis for comparison with the mammalian preBötzinger complex (Bouvier et al., 2010).

*SGNm-IRr* illustrates how activity-defined ensembles can refine the organization of populations previously implicated in behavior. Strong *c-fos* labeling in *calb2a*-expressing neurons of the secondary gustatory nucleus (SGN) after prey hunting led to the proposal that these neurons respond to food ingestion (Shainer et al., 2023). Our analysis distinguished medial and lateral SGN populations, with the medial population participating in a distributed ensemble with the inferior raphe (**Figs. 3D,E** and **6**). This ensemble was active in the absence of prey and outside episodes of eye convergence associated with hunting, suggesting spontaneous activity within feeding-related circuitry or a broader role in gustatory, visceral, or internal-state processing. Such functional diversity is compatible with the topographically organized gustatory and visceral inputs described in the adult zebrafish SGN (Yáñez et al., 2017). Within *SGNm-IRr*, independent expression-map and same-cell WARP analyses identified shared *calb2a* enrichment in the medial SGN and inferior-raphe populations, while registered neuronal reconstructions revealed candidate projections in both directions between their territories ((Du et al., 2026; Marquez-Legorreta et al., 2026; Shainer et al., 2023; Song et al., 2025); **Fig. 6D,E**). These observations motivate a model in which recurrent interactions contribute to the ensemble’s distributed activity. Calretinin could additionally regulate neuronal excitability and firing-rate gain through its calcium-buffering properties (Faas et al., 2007; Gall et al., 2003), although its contribution to *SGNm-IRr* dynamics remains untested. Simultaneous recordings from medial and lateral SGN and the inferior raphe during prey detection, ingestion, and prey-free periods could distinguish feeding-related recruitment from broader ongoing activity, while targeted perturbations and synaptic reconstruction could test the proposed recurrent organization.

The proposed correspondences of SGN with the parabrachial complex and rostral inferior raphe with raphe magnus suggest a mammalian comparison (Song et al., 2025; Volkmann et al., 2010). In mammals, parabrachial inputs reach neurons in the rostral ventromedial medulla, and Tacr1-expressing RVM neurons project back to the lateral parabrachial nucleus (Chen et al., 2017; Prajapati et al., 2025). mapzebrain expression maps also suggest *tacr1a* expression in the zebrafish inferior-raphe territory, providing a candidate molecular link. Whether these receptor-expressing cells contribute to SGNm-IRr, and whether the regional similarity across species extends to cell type and function, remains to be tested.

An important next step is to determine whether these co-activity patterns persist across developmental stages and are shared with other species. Comparisons across zebrafish development could reveal how distributed ensemble organization changes with maturation, while other fish models, such as *Danionella*, allow corresponding organization to be examined in an optically accessible adult nervous system (Schulze et al., 2018). Beyond fish, the proposed mammalian counterparts compiled in **Supplementary Table 1** provide a basis for testing whether co-activity among anatomically distant hindbrain populations is conserved. Whole-hindbrain experiments in zebrafish generate specific hypotheses about which mammalian populations are co-active and under which sensory or behavioral conditions. These predictions could guide the selection of populations for simultaneous recording in the mammalian brainstem, where whole-hindbrain cellular-resolution recording is not currently feasible (Agetsuma et al., 2025). Such targeted experiments would test whether conservation extends beyond the identity of individual populations to their organization into distributed functional ensembles.

The atlas captures reproducible co-activity under the conditions sampled but is not exhaustive. Shared-variance decomposition favors populations with sufficient co-active signal, and our filtering explicitly excluded small or weakly represented groups. Limited light-field resolution further constrains the separation of adjacent populations, particularly in the most ventral and dorsal regions. Recordings also did not consistently extend to the caudalmost hindbrain, including the area postrema, so populations in this region may be underrepresented. Beyond these detection limits, atlas coverage also depends on the sensory and behavioral conditions sampled. Our recordings were restricted to head-fixed larvae at 5 days post-fertilization, a preparation that removes the vestibular stimulation normally generated by body motion and alters mechanosensory feedback. We also sampled a limited behavioral repertoire, notably excluding hunting and escape responses. Networks preferentially recruited by these missing inputs or behaviors may therefore be underrepresented or missed. For the ensembles recovered, incomplete behavioral characterization means that low in-sample explained variance does not establish independence from behavior; even where coupling is clear, calcium-response kinetics constrain the temporal precision of activity–behavior relationships. Linking these ensembles to their underlying cells and circuits requires similar caution. Spatial overlap with expression maps from different animals nominates candidate molecular identities but does not establish that the participating neurons express the corresponding markers. This distinction could only be resolved for neurons contributing to matched ensembles in the WARP dataset, where activity and gene expression were measured in the same cells (Marquez-Legorreta et al., 2026). Likewise, co-activity and overlapping neuronal projections nominate candidate circuit relationships without establishing direct synaptic connectivity.

Our study identifies hindbrain ensembles that recur across individuals, with a core set robustly recovered across independent datasets, reporter lines, imaging modalities, and analysis methods. These ensembles show strong bilateral organization, spatial correspondence to gene-expression patterns, and different activity timings relative to movement. The atlas also provides testable hypotheses about functional coupling among the putative homologues of co-active zebrafish populations, particularly in vertebrates where simultaneous whole-hindbrain activity mapping at high spatial resolution is not yet possible.

## Data and code availability

All experimental and simulation data, Python scripts for data analyses and simulations, as well as to generate plots, are available at the KonData repository with the persistent identifier https://doi.org/10.48606/xf02th13j8jc6vd6. We will also seek to make the anatomical and functional masks available in the main atlases and repositories. Requests for further information and resources should be directed to and will be fulfilled by Armin Bahl. Source code implementing the algorithm for the light field deconvolution has been published previously (Aimon et al., 2019) and is available under https://github.com/sophie63/FlyLFM.

*The above-mentioned persistent identifier will only work upon final paper acceptance. For the review process, please use the following temporary link containing the data and source code archives:* https://cloud.uni-konstanz.de/index.php/s/kpe4BbPnEDFTyHm.

## Supporting information

Video S1. Functional light-field imaging in behaving larvae

Supplementary Table 1. Established zebrafish hindbrain functional populations: functions, mask provenance, and proposed mammalian counterparts.

Table 1. Per-ensemble summary of the curated hindbrain ensembles.

## Acknowledgments

We would like to heartily thank Liangyu Tao, Panagiotis Eleftheriadis, Margherita Zaupa, Meha Jadhav, Daniel Hummel and Sydney Hunt for providing helpful feedback on the manuscript. Alice Wang and the Bahl laboratory members for helpful discussion. Einat Couzin-Fuchs generously lent the backbone of the microscope and the Teledyne camera. The authors would further like to thank the state of Baden-Württemberg for its support of bwHPC and the German Research Foundation (DFG) for funding under “Project number 455622343” (bwForCluster NEMO2; compute project bw23A005).

## Author contributions

Conceptualization: S.A., A.B.; Methodology: S.A., A.B.; Software: S.A., A.B.; Validation: S.A.; Formal analysis: S.A.; Investigation: S.A.; Resources: A.B., H.N.; Data Curation: S.A.; Writing - Original Draft: S.A.; Writing - Review & Editing: S.A., A.B., H.N.; Visualization: S.A.; Supervision: A.B.; Project administration: A.B.; Funding acquisition: A.B.

## Declaration of generative AI and AI-assisted technologies in the manuscript preparation process

During the preparation of this work, the authors used ChatGPT (versions GPT-4 to GPT-6, OpenAI) and Claude Opus 5 (Anthropic) to assist with literature searches and synthesis, manuscript organization, and language editing. The authors independently consulted the original literature, verified all references and factual claims, and critically reviewed and revised all AI-assisted text. Scientific interpretations and conclusions were determined by the authors, who take full responsibility for the content of the article. The use of AI-assisted tools for code development is described in the STAR★Methods. All illustrations, such as the setup image in **Fig. 2A**, were designed and drawn by the authors manually or with custom-written Python code.

## STAR★Methods

### KEY RESOURCES TABLE

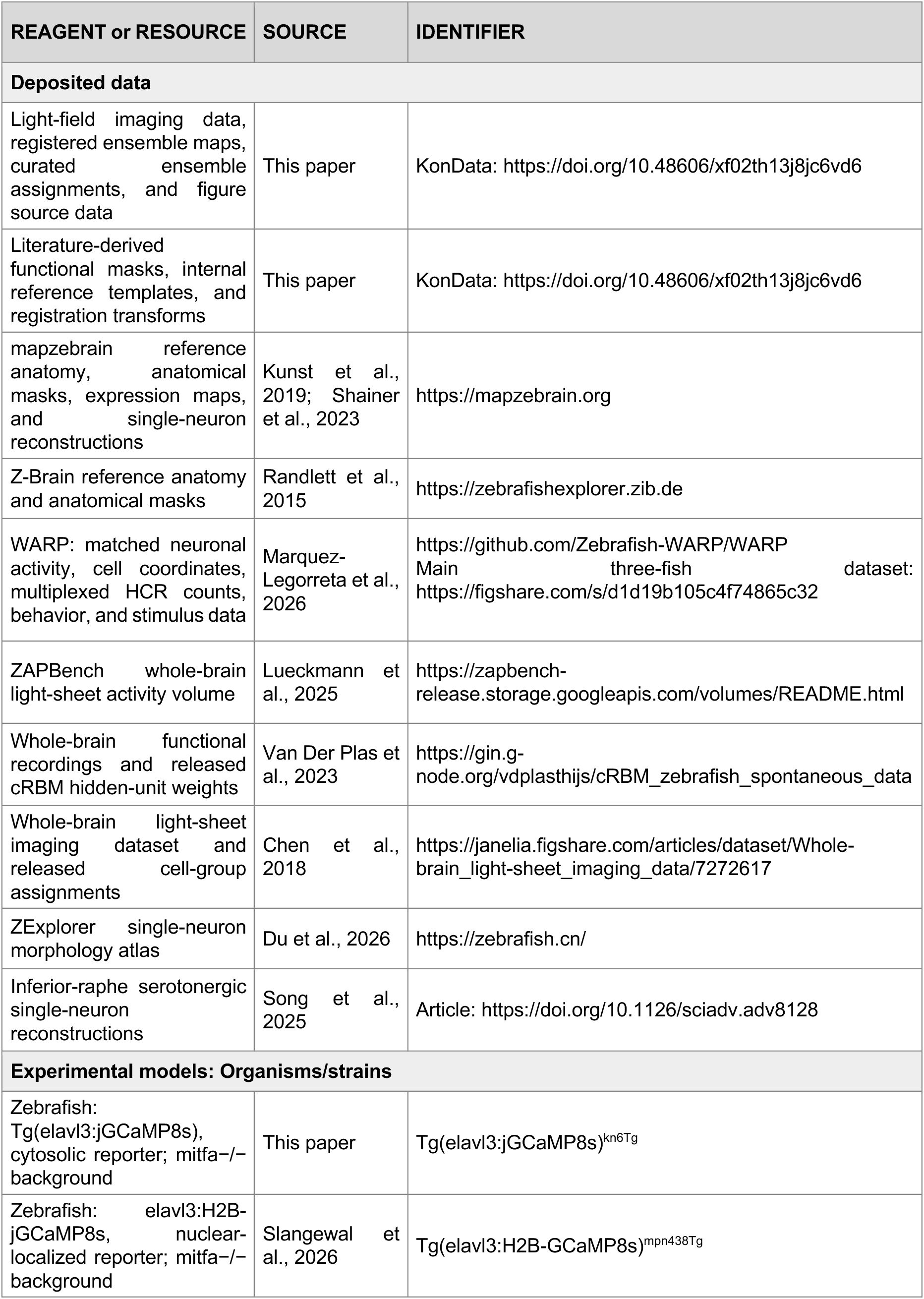
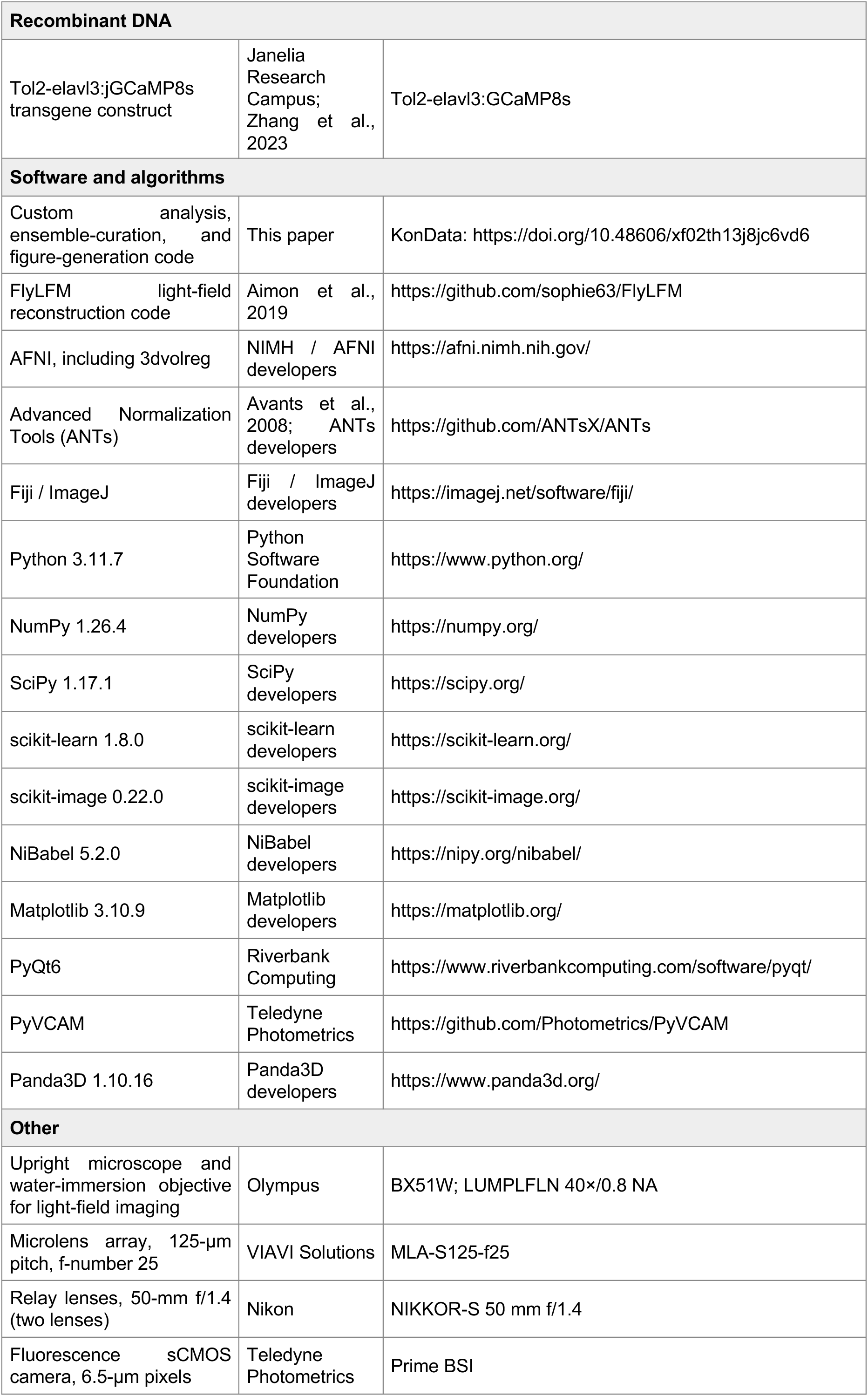
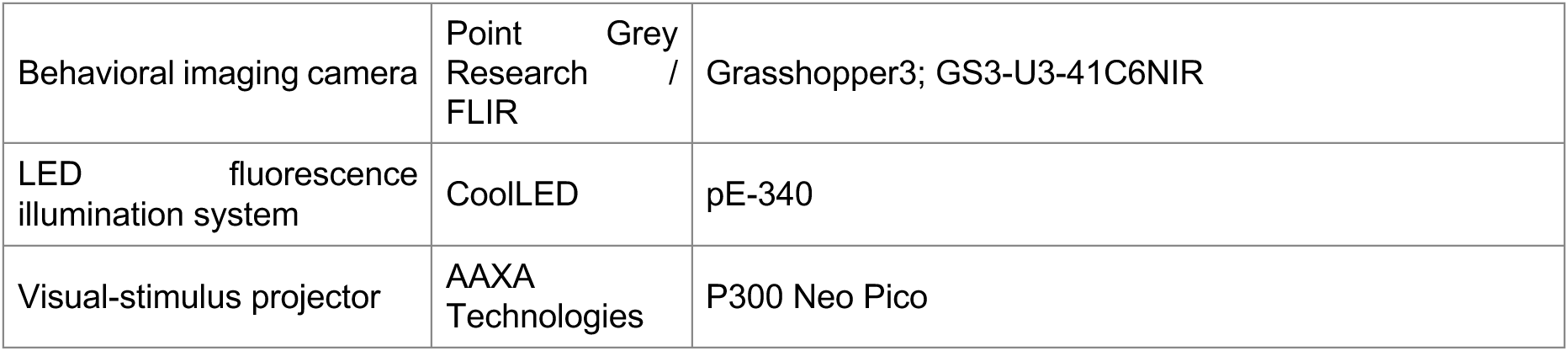

### RESOURCE AVAILABILITY

#### Lead contact

Further information and requests for resources should be directed to the lead contacts, Sophie Aimon or Armin Bahl.

#### Materials availability

We will share the newly generated cytosolic *elavl3:jGCaMP8s* line upon request. No additional reagents were created in this study.

#### Data and code availability

The light-field imaging data, registered ensemble maps, curated ensemble assignments, figure source data have been deposited at the KonData repository with the persistent identifier https://doi.org/10.48606/xf02th13j8jc6vd6. The repository also contains all source code to generate figures from the original preprocessed datasets. Public datasets re-analyzed in this study were obtained from (Chen et al., 2018; Lueckmann et al., 2025; Marquez-Legorreta et al., 2026; Van Der Plas et al., 2023).

### EXPERIMENTAL MODEL AND SUBJECT DETAILS

#### Zebrafish lines, husbandry, and ethics

We generated the Tg(*elavl3:jGCaMP8s*)*^kn6Tg^* line, which drives broad neuronal expression of jGCaMP8s in the cytosol. To establish the line, we co-injected 1–4-cell-stage embryos with the Tol2-elavl3:jGCaMP8s transgene construct (Zhang et al., 2023), provided by Janelia Research Campus, and Tol2 transposase mRNA. Injected founders were raised and crossed, and stable transgenic carriers were selected in the following generation based on strong green fluorescence throughout the central nervous system. Experimental larvae were obtained by incrossing these carriers.

Experiments used 5-days-post-fertilization larval zebrafish expressing pan-neuronal jGCaMP8s either in the cytoplasm (Tg(elavl3:jGCaMP8s)^kn6Tg^ , generated in this study) or in nuclei (Tg(elavl3:H2B-GCaMP8s)^mpn438Tg^; (Slangewal et al., 2026; Zhang et al., 2023)). All fish were in the *mitfa^−/−^* background to reduce pigmentation for the imaging. Our dataset consists of 68 larvae imaged in 84 whole-hindbrain recordings: 45 larvae expressing nuclear-localized jGCaMP8s (50 recordings) and 23 expressing cytosolic jGCaMP8s (34 recordings). Individual larvae could contribute more than one recording; analysis-specific recording counts and inclusion criteria are given below. All experiments were done in accordance with holding approval by the Regierungspräsidium Freiburg, Germany.

### METHOD DETAILS

#### Literature-derived atlas of hindbrain functional populations

**Fig. 1** was assembled in mapzebrain atlas space (Kunst et al., 2019; Shainer et al., 2023), and the complete mask set was also transformed to Z-Brain space (Randlett et al., 2015). Existing atlas parcels were used directly where they corresponded to a published functional population. Other masks were constructed by combining or subdividing atlas parcels, transferring registered functional maps, manually delineating regions in Fiji or python GUIs from published localizations, or thresholding registered expression maps. **Supplementary Table 1** documents the source and construction of each mask. Masks were organized into eight non-exclusive functional groups: eye control, locomotion, vestibular, mechanosensation, neuromodulation, orientation integration, orofacial–visceral, and cerebellum-associated. These groups organize the atlas without implying functional isolation between systems.

Eye-control masks included the oculomotor, trochlear, and abducens nuclei from mapzebrain. The velocity-to-position neural integrator was represented by an operational region of interest in medial caudal rhombomeres 7–8, guided by published functional and anatomical localizations (Brysch et al., 2019; Lee et al., 2015; Miri et al., 2011). The saccadic burst-generator ROI was manually delineated in medial rhombomeres 5–6 using published premotor localizations (Brysch et al., 2019; Dowell et al., 2025; Schoonheim et al., 2010).

Locomotor masks combined several sources. The nucleus of the medial longitudinal fasciculus (nMLF) mask was taken directly from mapzebrain, and the Mauthner, MiD2cm, and MiD3cm masks were combined from Z-Brain. The forward-swim mask combined Z-Brain RoL/RoR, MiR, MiT, and MiV parcels, guided by the functional organization described by (Lau et al., 2025). The strong-turn mask was generated by thresholding the mapzebrain *vsx2* reporter map (nns18Tg), informed by studies of V2a reticulospinal neurons (Collins et al., 2025; Kimura et al., 2013). The J-turn ensemble mask was a composite ROI based on published locations of MeLc/MeLr neurons and lateral reticulospinal populations (Gahtan et al., 2005; Lau et al., 2025). The mesencephalic locomotor region and prepontine long-latency C-start masks followed published localizations (Carbo-Tano et al., 2023; Marquart et al., 2019). These masks delineate spatial populations without assigning a separate mask to each activity pattern described by (Lau et al., 2025).

Vestibular masks comprised the octaval ganglion, superior, medial, and tangential vestibular nuclei, and vestibulospinal population. Z-Brain masks were used for the octaval ganglion and medial and tangential vestibular nuclei. A coarse superior-vestibular ROI was manually delineated from published localizations (Jia and Bagnall, 2022; Liu et al., 2022; Straka and Baker, 2013). The vestibulospinal ROI followed spinal-backfill and vestibulospinal-neuron localizations (Liu et al., 2022, 2020). The mechanosensory group included the posterior lateral-line ganglion and medial octavolateralis nucleus, represented by existing atlas masks.

Neuromodulatory masks comprised the locus coeruleus, superior and inferior raphe, nucleus isthmi, and medullary catecholaminergic groups. Population assignments followed published transmitter and anatomical characterizations (Filippi et al., 2010; Guo et al., 1999; Henriques, 2019; Holzschuh et al., 2003; Kastenhuber et al., 2010; Lillesaar et al., 2009). The superior- and inferior-raphe atlas masks were retained as parent ROIs. **Supplementary Table 1** relates these territories to the rostrocaudal SR-R, SR-M, SR-C, IR-R, and IR-C subsystems described by (Song et al., 2025).

Orientation-integration masks included the interpeduncular nucleus, heading-direction neurons, hindbrain oscillator/anterior rhombencephalic turning region (HBO/ARTR), anterior-hindbrain motion integrator, motion-onset/direction-selective population, and nucleus incertus. The mapzebrain interpeduncular mask was divided into dorsal and ventral regions using published anatomical descriptions (Aizawa et al., 2011, 2005; Bianco and Wilson, 2009). Heading-direction and HBO/ARTR ROIs followed published localizations (Dunn et al., 2016; Petrucco et al., 2023; Wolf et al., 2017). Motion-integrating and motion-onset ROIs were transferred from registered functional maps (Bahl and Engert, 2020; Slangewal et al., 2026). Nucleus incertus subregions associated with *gsc2* and *rln3a* were manually delineated following (Spikol et al., 2024).

Orofacial–visceral masks included the trigeminal, facial, and vagal motor nuclei, vagal sensory lobe, and area postrema from mapzebrain. The glossopharyngeal motor ROI was manually delineated around paired lateral *chata*/*isl1*-positive somata between the facial and vagal motor masks, guided by published branchiomotor anatomy (Chandrasekhar, 2004; Chandrasekhar et al., 1997; Mueller et al., 2004). Additional operational ROIs were manually drawn to approximate the gut nutrient-responsive dorsal vagal complex and medial idMO territories reported by (Chen et al., 2026). The secondary gustatory/viscerosensory nucleus ROI was generated by smoothing and intensity-thresholding the local isthmic *calb2a* expression domain, using published larval localization to guide its assignment (Henriques, 2019; Shainer et al., 2023).

Cerebellum-associated masks comprised the cerebellar cortex, eminentia granularis, and inferior olive, with population annotations informed by published anatomical and functional studies (Bae et al., 2009; Félix et al., 2024; Kani et al., 2010; Knogler et al., 2019; Takeuchi et al., 2015).

The reference masks primarily delineate neuronal cell-body territories, although the interpeduncular nucleus and some cerebellar masks also encompass neuropil. Masks may overlap where their definitions share anatomical territory. Populations without a localized mask, including the respiratory rhythm generator, were retained as reference entries in **Supplementary Table 1** but were not represented spatially in **Fig. 1**.

#### Cross-species homology assignment

Each zebrafish population was assigned a putative mouse counterpart by integrating rhombomeric position and Hox identity, molecular phenotype, connectivity, physiological function, and developmental lineage, where available ((Di Bonito et al., 2015; Straka and Baker, 2013) **Supplementary Table 1**). Confidence tiers were assigned qualitatively based on the combination of evidence summarized in **Supplementary Table 1**; no numerical scoring scheme was applied. Assignments were designated strong when developmental ancestry supported the functional and positional correspondence; moderate when functional, molecular, or positional evidence was substantial but shared lineage was unconfirmed or development diverged; and open/weak when the correspondence was tentative, or no direct mammalian homologue was known. Mouse structures were illustrated schematically in **Fig. S1**, using the same functional categories and population colors as **Fig. 1**. These schematics were not registered to a mouse atlas and indicate approximate positions only; dot size and spacing do not represent anatomical size or spatial relationships. **Supplementary Table 1** provides the evidence, confidence tier, and references for each assignment.

#### Light-field imaging and behavioral acquisition

Whole-hindbrain calcium activity was recorded in head-fixed larval zebrafish by light-field microscopy, which encodes information from the imaged volume in a single camera exposure (Aimon et al., 2023, 2019; Broxton et al., 2013). Larvae were imaged at 5 days post-fertilization, with movements of the eyes, pectoral fins, tail, and mouth available for behavioral observation.

The fluorescence optical arrangement followed Aimon et al. (2019). An upright Olympus BX51W microscope was equipped with an Olympus LUMPLFLN 40×/0.8-NA water-immersion objective and a VIAVI Solutions microlens array with a 125-µm pitch and f-number of 25. The array was positioned at the microscope image plane, and its back focal plane was relayed onto the camera sensor using two Nikon 50-mm f/1.4 NIKKOR-S lenses. We used a CoolLED pE-340 LED system light source and a GFP filter cube. Fluorescence was detected with a Teledyne Photometrics Prime BSI sCMOS camera with 6.5-µm pixels.

Exposure and sampling rate were selected for each recording. Most cytosolic-reporter recordings used 200- or 50-ms exposures, corresponding to approximately 5 or 20 Hz, respectively; most H2B-reporter recordings used 200-ms exposures.

Fluorescence acquisition, behavioral recording, and visual stimulation were coordinated by custom Python software. A PyQt6 interface controlled acquisition, tracking, stimulus, and experiment-control processes. The fluorescence camera was operated continuously with internal triggering through the PyVCAM/PVCAM interface. Images were stored as unsigned 16-bit arrays in HDF5, together with beginning-of-frame camera timestamps. The acquisition configuration and a copy of the selected stimulus script were saved with each experiment.

Behavior was recorded simultaneously with a Grasshopper3 GS3-U3-41C6NIR camera (Point Grey Research/FLIR) under infrared illumination, at approximately 178 Hz. Camera exposures were 3–5 ms, and square acquisition regions of interest ranged from 256 × 256 to 736 × 736 pixels, depending on the recording. The tail midline was extracted online from an operator-defined tail base. Successive nodes were located by sampling image intensity along an arc extending ±60° around the previous segment direction, in 2° increments, smoothing the angular intensity profile and following its darkest direction. Tail-tip deflection was calculated from the mean angle of the final five nodes relative to the tail base. Behavioral videos, frame timestamps, and numerical tracking results were saved for subsequent analysis of tail, eye, fin, and mouth movements.

Visual stimuli were presented using a P300 Neo Pico projector (AAXA Technologies). Visual stimuli were rendered using Panda3D at a target update rate of 60 Hz, with the display configured to use the red channel. Conditions included a black display, textured visual scenes, and drifting sinusoidal gratings. In naturalistic closed-loop trials, a periodically tiled background image (a rice-paddy scene), was translated and rotated in response to estimated swimming movements. Earlier textured-scene protocols updated the scene following detected swim bouts. Stimulus identities and onset and offset times were recorded with the experiment.

For closed-loop trials using a 5-dpf autoregressive tail decoder, the tracked tail was translated to its base, aligned to its first segment, resampled to 30 equally spaced points along its length, and normalized by total tail arc length. Differences between consecutive normalized midlines provided 60 standardized features. A ridge-regression model with fitted intercepts estimated forward displacement, lateral displacement, and heading change from the current and 40 preceding tail-feature vectors and 15 preceding predictions of the corresponding output (ridge penalty, 10⁻⁵; training target offset, six retained tracking samples). The model was trained using paired tail-posture changes and frame-to-frame movements of freely swimming wild-type larvae at 5 days post fertilization, recorded at a nominal 178 Hz. Fourteen fish with valid tail midline tracking contributed 429,136 samples to fitting the feature scaler. Each regression used 2,475 inputs: 41 vectors of 60 tail features and 15 values from the history of its corresponding output. Training used measured output histories, whereas online decoding used previous predictions. Decoder output was gated by a recording-specific tail-motion threshold and used to update the visual texture, with scene translation and rotation opposing estimated animal movement.

Temporal information was retained separately for the fluorescence, behavioral, and stimulus streams. Fluorescence timestamps were obtained from camera beginning-of-frame metadata, and behavioral timestamps from the FLIR camera clock.

#### Volume reconstruction, preprocessing and registration

Raw light fields were reconstructed by volumetric deconvolution into three-dimensional image stacks with 3-µm isotropic voxels (Aimon et al., 2019; Broxton et al., 2013). Source code implementing the algorithm is available under https://github.com/sophie63/FlyLFM. Reconstructed volumes were motion-corrected by rigid-body (six-parameter) volumetric registration to the recording’s temporal-average volume using AFNI 3dvolreg (single-pass, one-voxel zero-padding). Frames failing a local-correlation quality check (median-absolute-deviation z > 3.5) were additionally re-registered to a mean of the passing frames with ANTs (rigid + affine + SyN). ΔF/F was then computed as (F − F0)/(F0 + 1e-6), where F0 is a centered 500-frame moving average (uniform/boxcar mean over time; 100 s at the 5 Hz imaging rate) of the brain-masked, motion-corrected stack.

Whole-hindbrain maps were registered to an internal symmetric reference template, defined in the light-field reconstruction grid (150 × 150 × 76 voxels). To build the template, the time-averaged reconstructed volumes of approximately ten larvae were each registered to a single initial reference brain with ANTs (rigid, then affine, then a symmetric normalization (SyN) nonlinear warp, cross-correlation similarity metric), and the aligned volumes were combined into one volume by distance-weighted feathered blending: at each voxel, every contributing volume was weighted by its Euclidean distance from its own tissue boundary; the weights were smoothed with a Gaussian (≈2.5 voxels) and normalized across volumes. The combined volume was made left–right symmetric by averaging it with its reflection across the midline. Every recording of both reporter lines was registered to this symmetric template with the same ANTs sequence, and each independent-component map was carried into the reference frame by the resulting transform; cytosolic-line recordings were additionally aligned to the nuclear-localized template by a thin-plate-spline landmark warp.

For each recording, a mean anatomical volume was formed as the temporal average of the motion-corrected reconstructed stack and, after a manually chosen rotation, registered with ANTs to a left–right–symmetric light-field reference template (Avants et al., 2008). Registration used a multi-resolution rigid → affine → deformable (symmetric normalization, SyN) cascade. The similarity metric was mutual information with 32 histogram bins for the rigid stage and neighborhood cross-correlation with a radius of 4 voxels for the affine and SyN stages; intensity histogram matching was disabled. Transform gradient steps were 0.1 for the rigid and affine stages and 0.1 for SyN, with SyN update- and total-field smoothing of 6 and 1 voxels. The rigid and affine stages were optimized over a five-level Gaussian pyramid (shrink factors 12, 8, 4, 2, 1; smoothing 12, 8, 4, 2, 2 voxels) and the SyN stage over a four-level pyramid (shrink factors 8, 4, 2, 1; smoothing 8, 4, 2, 1 voxels), with a convergence threshold of 1×10⁻⁸. The estimated forward warp, together with its inverse, was saved and applied to that recording’s ensemble maps to carry all functional data into the reference frame.

Cytosolic-line recordings were aligned in two steps. Each cytosolic recording’s mean anatomical volume was first registered, with the same ANTs cascade (SyN gradient step 0.05), to a separate left–right–symmetric template built from the cytosolic recordings; that cytosolic template was then aligned to the H2B light-field reference by a thin-plate-spline transform fitted to 24 manually placed anatomical landmarks. Combining the two transforms placed cytosolic ensembles in the same reference frame as the H2B line.

Z-Brain (Elavl3-H2B-RFP template channel) and mapzebrain (HuC/D channel) were brought into the same light-field reference. The mapzebrain reference was registered to the light-field reference by a thin-plate-spline transform fitted to 33 manually placed anatomical landmarks. Z-Brain was first aligned to mapZeBrain2024 with an ANTs deformation field and then carried into the light-field frame by the same thin-plate spline; anatomical parcels and expression maps were resampled through these transforms. Functional ensembles, cluster averages, anatomical parcels, and expression maps used for quantitative comparisons therefore occupied a common 150 × 150 × 76 voxel grid.

#### Spatial ICA and ensemble filtering

Each recording was decomposed by principal component analysis followed by independent component analysis on the principal component maps, into approximately 2,500 components, each consisting of a three-dimensional spatial map and an associated time series (Aimon et al., 2019; Beckmann and Smith, 2004; Mukamel et al., 2009). No spatial-compactness constraint was imposed.

Components were filtered using prespecified spatial criteria designed to retain population-scale (ensemble) signals and exclude isolated compact sources. In particular, the requirement for at least two spatially separated peaks excluded isolated objects that, especially in cytosolic recordings, could not be reliably distinguished from blood cells and, in either reporter line, could represent single neurons or very small neuronal populations rather than the ensembles targeted here. Components were retained when they satisfied three criteria on the ANTs-registered map after 3-D Gaussian smoothing (σ = 2.5 voxels) and restriction to an analysis mask formed from the eroded brain mask (3 erosion iterations), with blood vessels excluded and the result intersected with a hindbrain mask. First, at least 50 voxels survived a selection threshold of 10·σ_noise, where σ_noise is the standard deviation of the non-zero values of the smoothed, masked maximum projections of each recording’s 200 lowest-ranked components. Second, at least two maxima on the log-transformed z-projection exceeded 0.6·σ_peak (σ_peak estimated from the log-transformed projections of the same 200 components), with each secondary peak at least 1% of the primary-peak height and a minimum peak separation of 6% of the projection’s linear dimension (maps were capped at 10 peaks). Third, the summed signed intensity inside the mask exceeded 50% of the summed signed intensity over the whole map. Filtering retained approximately 300 ensembles per recording.

The atlas-building dataset comprised 84 recordings from 68 larvae (nuclear-localized line, 50 recordings from 45 larvae; cytosolic line, 34 recordings from 23 larvae) and yielded 24,440 filtered ensembles. Curation (below) assigned 8,450 of these to 79 non-junk ensembles; the reproducible-ensemble atlas analyzed in **Figs. 2–6** was the subset meeting the membership criteria defined in the next section (70 clusters). Analysis-specific pools differed where additional recordings or uncurated filtered ensembles were required, as described below.

#### Cluster average maps and noise normalization

Each ensemble map was expressed in units of local noise: in native imaging space, it was divided voxel by voxel by its recording’s noise map (the per-voxel standard deviation across the 500 lowest-ranked component maps, median-smoothed with a 5-voxel kernel and floored at the 10th percentile of its non-zero values), and the result was registered to the reference with the same transform as the ensemble maps. Each ensemble’s average map was the mean of these noise-normalized member maps in the common left–right orientation described below, retaining only voxels where the mean magnitude exceeded 4/√n in noise units (n = contributing ensembles). For display, average maps used a hard-zero baseline with signed gray (positive) / magenta (negative) rendering, per-map normalized to the 99th percentile of projected magnitude, with a display gamma of 0.75.

#### Cross-recording feature extraction and ensemble clustering

For cross-recording clustering, lateralized ensemble maps were placed in a common orientation: maps with greater total loading in the right hemisphere were mirrored so that the dominant side was always on the left. This prevented left- and right-sided instances of the same spatial pattern from being assigned to separate clusters. We calculated 16-bin marginal loading profiles along the medial– lateral, anterior–posterior, and dorsoventral axes, standardized the marginal profiles, and combined them with a standardized log-transformed measure of the number of spatially distant peaks. Sixteen-bin marginal loading profiles were computed along three axes (dorsoventral depth, medial–lateral, and anterior–posterior, the two horizontal marginals in bilateral orientation). Each marginal was z-scored within ensemble and then z-scored across ensembles, giving a 48-column block. The peak-number term was log(1 + n_distant_peaks), using the same z-projection peak detector as the filtering step; this single column was z-scored across ensembles and multiplied by 3.0, so it carried three times the standard deviation of any individual marginal bin. The final feature vector had 49 columns.

A Gaussian-mixture model on the 49-column feature matrix (100 mixture ensembles, full covariance, n_init = 5, max_iter = 300, random_state = 0) provided the initial grouping, which was manually refined in a custom curation interface using registered maps and projection previews.

Spatially corresponding ensembles from different recordings were assigned to a common cluster. Each cluster was represented by the average spatial map of its member ensembles after consistent orientation. The reproducible atlas shown in **Fig. 2E** comprises 70 curated clusters with more than 10 member ensembles and contributions from at least 10 distinct larvae, with medians of 72 ensembles and 35 contributing larvae per cluster. t-SNE was calculated from the same feature matrix for visualization only and did not determine the final grouping (Van der Maaten and Hinton, 2008).

Out-of-sample reproducibility and grouping of ensembles. We quantified how reliably ensembles are recovered on new animals by leave-one-fish-out cross-validation. For each held-out fish, a linear discriminant classifier was trained on the ensembles of all remaining fish to assign each ensemble to one of the curated clusters. Each ensemble was described by the clustering feature vector (the 49-column standardized marginal-plus-log-peak representation) concatenated with a projection-on-average block: the Pearson correlation of the ensemble’s noise-normalized map in this common left– right orientation—spatially downsampled eightfold, averaging each 8 × 8 × 8-voxel neighborhood into one 24-µm voxel—to each of the 70 cluster averages, each ensemble average rebuilt from the training larvae only. The classifier was a linear discriminant analysis evaluated leave-one-fish-out over the 70-ensemble **Fig. 2E** set (8,384 ensembles from 62 larvae), with confidence intervals from a 2,000-sample bootstrap over fish. Crucially, these ensemble averages were recomputed from the training fish only, excluding the held-out fish, so no information from a test animal entered its own features or the classifier. Performance was summarized as balanced accuracy (the mean across ensembles of the fraction of each ensemble’s held-out ensembles assigned to it, weighting every ensemble equally regardless of size) and macro-F1; 95% confidence intervals were obtained by bootstrapping over fish (2,000 resamples). For display in **Fig. 2E**, ensembles were grouped by mutual out-of-sample confusability: two ensembles were joined when each misclassified at least 15% of its held-out ensembles as the other, and groups are the connected ensembles of these links, so an ensemble recovered below threshold on its own but sharing no strong reciprocal confusion with another is shown separately. The held-out assignments and the resulting grouping were produced automatically by the classifier; the manual curation contributed only the ensemble labels used as training targets, and no manual judgment entered the cross-validated assignments or the grouping.

#### Ensemble nomenclature and anatomical annotation

Descriptive ensemble names summarize the strongest available anatomical or molecular correspondence and do not imply exact identity with a previously defined nucleus, molecular cell type or circuit. Candidate atlas regions, anatomical and molecularly defined, were ranked by the fraction of the ensemble’s supra-threshold footprint mass falling within each region (computed on the SNR-thresholded ensemble average maps), excluding generic container regions; the highest-ranked region was used. Figure annotations combined manually assigned names with atlas regions ranked by enrichment.

#### Validation in independent functional datasets

We tested recovery of the curated ensembles in four independent whole-brain light-sheet datasets that differed in imaging modality, spatial representation and decomposition method: spatial PCA–ICA applied to segmented cells from the WARP recordings of (Marquez-Legorreta et al., 2026) (three larvae from the main release and three pilot larvae); spatial PCA–ICA applied voxel-wise to one ZapBench recording (Lueckmann et al., 2025); the authors’ released compositional restricted Boltzmann machine assemblies from seven recordings (Van Der Plas et al., 2023); and the authors’ released k-means correlation clusters (k = 20) from 15 subjects (Chen et al., 2018). Three Chen et al. subjects were excluded because the released archives could not be decompressed, and one cRBM recording was excluded because its released neuron subset did not match the weight matrix.

The external datasets used nuclear-localized or cell/voxel representations that did not reproduce neuropil-rich cytosolic ensembles. Ensembles for which at least 80% of curated members came from the cytosolic reporter were therefore designated cytosolic-line-dominated. Eleven such ensembles belonged to the 70-ensemble **Fig. 2E** set and were excluded before cross-dataset recovery was scored, leaving 59 eligible ensembles. These eleven were dominated (≥80% of members) by the cytosolic jGCaMP8s line, whose cell- or voxel-based representations do not reproduce neuropil-rich cytosolic ensembles.

#### Ensemble extraction from external datasets

For WARP, hindbrain and cerebellar cells were selected from the aligned segmentation. DF/F traces were concatenated within each recording and z-scored per cell over time. Each larva was decomposed into 2,500 spatially independent components. Whitening was performed in two passes: a first principal component analysis estimated a per-cell residual noise level (the standard deviation of the residual after removing the sparsified leading principal components), each cell was divided by this level; and a second principal component analysis on the noise-normalized matrix produced the components fed to FastICA (parallel algorithm, cube nonlinearity, tolerance 1e-3, <=1000 iterations). Scan-geometry “seam” components—thin, high-amplitude ridges spanning an imaging plane—were identified by a line-filter ridge score (peak-minus-flank profile amplitude, normalized by total signal) exceeding 0.03 in either the dorsal or sagittal raster of the absolute loading map; the temporal subspace of the flagged components was regressed out and the volume re-decomposed, iterating up to four times or until no component exceeded the threshold. Because spatial focality did not reliably separate biological from speckled components at this decomposition size, retained ensembles were selected by temporal coherence, requiring lag-1 autocorrelation >= 0.8. This yielded 185, 331, and 181 retained ensembles in the three main larvae and 139, 140, and 444 in the three pilot larvae.

The downsampled ZapBench volume was decomposed into 1,000 components with the identical two-pass whitening and cube-FastICA framework (seam components regressed out and the volume re-decomposed, as above). Stripe and scan-seam components were flagged by the same line-filter stripiness score exceeding 0.15 and removed (147 of 1,000), leaving 853 ensembles.

For cRBM data, released hidden-unit weight matrices were used directly. Hidden units whose maximum absolute weight fell below 10% of the per-recording median were discarded as inactive; within a recording, units sharing more than 60% of their top-weight member neurons (the neurons carrying 80% of a unit’s total absolute weight) were merged; and units with less than 30% of their absolute weight inside the hindbrain were excluded, leaving 631 hindbrain-enriched units (1,200 released −> 972 unique −> 631 hindbrain-enriched).

For the Chen et. al dataset, each released k = 20 cell grouping was rasterized into the reference grid by accumulating one count per cell at its rounded voxel and Gaussian-smoothing (sigma = 2, 2, 1 voxels along the mediolateral, rostrocaudal, and dorsoventral axes), giving a spatial density map.

#### Registration of maps from external datasets

The common chain was source space to Z-Brain, Z-Brain to mapzebrain by ANTs SyN, deterministic fourfold downsampling and cropping to a 110 × 100 × 25 working reference, and a landmark-based thin-plate-spline transform into the light-field reference. The complete chain was checked by propagating known atlas masks; the warped Z-Brain rhombencephalon had a Dice overlap of 0.60 with the light-field rhombencephalon. For segmented-cell data, regional centroids were compared after splitting bilateral regions into left and right halves, yielding residual errors of approximately 1.2–1.4 light-field voxels.

#### Spatial matching and recovery assignment

Curated ensemble averages and external dataset maps were compared by cosine similarity between focal spatial cores. For continuous maps, the 92nd percentile of the positive (non-zero) voxel values was subtracted, and negative residuals were clipped to zero, suppressing broad low-amplitude halos; binary density maps (segmented-cell methods) were compared without this subtraction. Before comparison, each map was left-right symmetrized (added to its mirror image across the mediolateral axis) so that a lateralized ensemble could match its mirror, and then unit-normalized. cRBM maps were additionally sign-flipped so that their strongest-magnitude lobe was positive.

Recovery provides a conservative, lower-bound measure of cross-method reproducibility and was not treated as a statistical test. Cosine similarity was used only to rank and pre-group candidate matches, with no cutoff excluding external dataset ensembles from curatorial review. Final identities were assigned manually in the same browser used to curate the light-field ensembles, with the corresponding ensemble average displayed as a reference.

For cell-based tiles in **Fig. S3**, member counts were restricted to the rhombencephalon. WARP members were cells with absolute ICA-loading z-score > 4.0. cRBM members were the largest-absolute-weight neurons carrying 85% of the unit’s total absolute weight. Chen counts included all rhombencephalic cells in the k-means group. Voxel-based methods were not assigned member-neuron counts.

#### Reference expression atlas and molecular decomposition

We assembled 747 registered mapzebrain expression maps, including transgenic and enhancer-trap reporter lines, HCR/ISH gene-probe stains, antibody stains, and dye labels. Each map was resampled into the common symmetric reference, restricted to the functional hindbrain mask and vectorized. The resulting voxel-by-map matrix was mean-centered and decomposed by PCA followed by ICA. The mean-centered 747-map voxel matrix was decomposed by FastICA (n_components = 746, algorithm = “parallel”, nonlinearity = “cube”, max_iter = 1000, tol = 1e-3) following a two-pass noise-normalizing whitening (a knee-selected first-pass PCA whose sparsified loadings, thresholded at 0.4·SD, were subtracted and the residual rescaled per voxel). Diffuse or noise-like components were identified from their supra-threshold spatial support: Each component map was divided by its standard deviation over the hindbrain and smoothed (σ = 3 voxels), and a component was flagged as diffuse when its largest connected component above 2.5 standard deviations of the smoothed map exceeded 20% of the hindbrain support. 88 components met this criterion and were excluded, leaving 658 sparse expression components for functional–expression matching and the subsequent molecular analyses. Each retained component represents an independently varying spatial expression pattern, whereas its loading vector gives the contribution of each input map. For the **Fig. 3B** overlay, 20 expression components were thresholded at a spatial-map value of 4 (components with at least 300 supra-threshold voxels) and shown as their whole-depth projection; the 20 anatomical regions displayed alongside were down-selected to reduce visual crowding (projected area at or below the 90th percentile, pairwise projected overlap at or below 0.10 of the smaller region), purely for legibility.

#### Functional–expression and functional–anatomical overlap

Functional ensemble averages, expression components, and anatomical regions were compared in the same 150 × 150 × 76 voxel grid. For quantitative overlap, the functional footprint was the unthresholded ensemble average added to its left–right mirror. Functional and expression maps were positive-clipped (negative voxels set to zero), atlas regions and tiles entered as binary masks, all maps were L2-normalized, and overlap was the cosine between normalized maps. For display in **Fig. 3E, F** and all overlays, functional average volumes were thresholded at 2.5 times the standard deviation of their non-zero voxels and normalized to their 99.5th percentile; expression maps were projected as absolute values, had the 10th percentile of their projected non-zero values subtracted and were normalized to their 99.5th percentile; both used a display gamma of 0.55. The **Fig. 3D** functional maps show the unthresholded average with the ensemble-map renderer (99th percentile, gamma 0.75). These steps did not affect the reported overlaps. The same preprocessing was applied to all candidate maps within a comparison.

For each of the 70 curated clusters, we calculated the maximum cosine overlap with any of the 178 atlas regions and with any of the 658 retained expression components. Atlas regions were the mapzebrain, Z-Brain and Fig. 1 masks, including molecularly defined clusters, stripes and neurotransmitter domains; masks sharing a region name (left and right) were merged into one bilateral mask, and generic containers were excluded (whole hindbrain, rhombomeres, medulla oblongata subdivisions, neuropil regions, cerebellum, corpus cerebelli and the cerebellar Vglut2-enriched areas). As an arbitrary-parcellation control, the left half of the rhombencephalon mask was partitioned by k-means on voxel coordinates into 658 contiguous tiles (matching the number of sparse expression components), and each tile was mirrored across the midline to form bilateral null blobs (matching the bilateral symmetry of the ensembles). For every functional ensemble, we took its maximum cosine overlap with these null blobs, through the same overlap pipeline. Because a null overlap magnitude has no meaningful anatomical partner, the shaded band in **Fig. 3C** spans the minimum-to-maximum of these per-ensemble null overlaps, drawn as a constant-width band along the anti-diagonal.

Featured single matches pair a functional ensemble with the retained component of highest cosine overlap minus that component’s mean overlap across all ensembles. Multi-component matches were built greedily (up to three components): the functional target was smoothed (3 µm), its broad pedestal removed by a white top-hat filter, restricted to voxels above 2.5 standard deviations and mirror-symmetrized; at each step the retained component with the highest baseline-corrected cosine to the remaining target was accepted if its territory (a hysteresis mask of the component smoothed at 10 µm, seeds at 35% and growth to 10% of its peak) removed at least 8% of the remaining target mass, and matching stopped when the best cosine fell below 0.18. To assign a molecular label we calculated the Pearson correlation between the top-down projection of each expression component and each of the 747 input-map projections reconstructed from the decomposition, over the brain support. DAPI, pan-neuronal, and indicator maps were excluded, as were broadly active maps occupying >55% of the brain; 379 input maps remained as interpretable candidate labels. The highest-correlated interpretable maps were reported.

The gray labels next to each ensemble name (**Fig. 3D**) and under each example (**Fig. 3E,F**) list the atlas regions, including molecularly defined masks, overlapped by the ensemble’s thresholded average map (the same SNR-thresholded ensemble volume shown in **Fig. 2E**), ranked by the fraction of the ensemble’s supra-threshold mass falling inside each region (regions contributing at least 8% are listed, up to four in **Fig. 3D** and five in **Fig. 3E,F**; the generic containers listed above are excluded). The black labels list the source expression maps most strongly correlated with the matched component, as described above.

#### Transmitter identity of spatial subclusters in WARP

The WARP recordings combined 2-Hz whole-brain functional imaging with multiplexed HCR measurements in the same segmented cells (Marquez-Legorreta et al., 2026). We analyzed three larvae from the main release (42-gene panel) and three pilot larvae (32-gene panel). Analyses used the hindbrain cells included in the functional decomposition from **Fig. S3** (61,030, 70,492 and 64,642 cells in the main larvae; 69,660, 66,577 and 68,516 in the pilot larvae) and their one-to-one indexed marker counts. Ensembles were extracted, filtered, registered, and assigned to curated ensembles as described above. For each displayed cluster, we included all ensembles manually assigned to it across the six larvae.

For each ensemble, the focal core comprised cells whose loading exceeded mean|w| + 5 SD|w| in either direction, with the same floor for positive and negative loadings. Because the registered cell centroids occupied a common template, cores from different larvae were pooled directly. Pooled cells were restricted to the functional hindbrain mask: the light-field mask (the summed absolute ensemble-average volume thresholded at 8% of its 99th percentile) was warped into the WARP template by an inverse thin-plate-spline landmark transform, and cell centroids were mapped by the landmark affine transform of one larva. This yielded 628–6,634 cells per ensemble in **Fig. S4** and 262–2,528 in **Fig. 3G**.

Each cell was assigned one transmitter class as the argmax of abundance-normalized marker counts, each marker divided by its mean across the decomposed hindbrain cells: *vglut2a*, *gad1b*, *glyt2*, *chata*, *tph2* and th in the main larvae, and *vglut2a*, *gad1b* and *tph2* in the pilot larvae; cholinergic identity was taken from the main larvae only. Assignment required at least two transmitter-marker spots in total and at least one count for the winning marker; cells failing these criteria were displayed as unassigned and excluded from percentage calculations. Across all decomposed hindbrain cells of the main larvae, the background fractions were approximately 26% glutamatergic, 20% GABAergic, 7% glycinergic, 7% cholinergic and 5% serotonergic, with approximately 35% unassigned or of a class not shown.

Within each ensemble, pooled focal-core cells were segmented into localized three-dimensional subclusters. Cells were binned and weighted by their ensemble loading; the density was Gaussian smoothed, local maxima seeded a watershed, and adjacent basins sharing the same dominant transmitter were merged; subclusters needed at least 20 cells. Detection used all core cells, and transmitter composition was calculated over typed cells only. In **Fig. S4**, a subcluster was outlined and labeled when one transmitter class constituted >50% of its typed cells and the subcluster held at least 10% of the ensemble’s typed cells; for *HBO*, the subclusters were re-divided at 40 template units from the midline into a medial part and a lateral part (left and right counted together), following the gap in its neurons’ mediolateral distribution. In **Fig. 3G**, the largest such subcluster of each class was outlined for glycinergic and cholinergic subclusters, the GABAergic subcluster of *VestLCRaph*e and the serotonergic subcluster of *SGNm*-*IRr*.

Gene-defined clusters (**Fig. 3G** and *npbCl* in **Fig. S4**, dashed outlines) were segmented in the same way from the density of gene-positive focal-core cells (at least two spots), without basin merging, in the larvae whose panel contains the gene. A cluster was kept when it held at least 8 gene-positive cells (5 for th) and its mean spot count was at least twice the hindbrain mean (pseudocount 0.02 spots per cell); all kept clusters of a gene share one label giving the fold enrichment over their pooled cells (for *VestLCRaphe* also of *net* and *uts1*, for *SGNm-IRr* also of *calb2a*), linked to each cluster by a leader line. Gene enrichments quoted in the text are the mean spot count of an ensemble’s focal-core cells relative to the hindbrain mean. The anatomical position of the Abd-Occ anterior cholinergic cluster was assigned from the light-field ensemble map, whose positive mass overlapped the atlas oculomotor nucleus 16-fold more than the trochlear nucleus.

For display, neurons were drawn from ventral to dorsal in the dorsal view and from the right to the left side in the depth view, so that the neurons nearest the viewer lie on top. The gray outline is the smoothed density of the hindbrain cells inside the functional mask, mirrored across the midline in the dorsal view.

#### Left–right symmetry classification

For the symmetry analysis (**Fig. 4**), the noise-normalized ensemble maps (see “Cluster average maps and noise normalization”) were used, expressed in units of local noise (20,060 ensembles across 68 recordings: H2B 39 recordings / 32 fish; cyto 29 recordings / 18 fish).

#### Symmetry classification

Functional and expression maps were classified by a single shared procedure. Each map was first thresholded at the noise level on the full-resolution volume, then binned to 24 µm for the symmetry measurements, just below the autocorrelation scale (we measured it to be 32 microns for expression components and 38 microns for functional ensembles). Because functional maps are already in noise units, a fixed cut of |value| >= 2.5 (i.e. 2.5 sigma) was applied; for expression, the noise sigma was estimated from the symmetric (negative-reflected) part of the value histogram of the five lowest-ranked components on the full-resolution masked maps, and the cut was 2.5 x sigma. Binning preserved the midline: the left and right halves of the x-axis were binned independently about the center, so a voxel and its mirror stayed aligned. Each component was then assigned a category in order:

1. Midline. A component was called midline if at least 50% of its thresholded absolute mass lay within +/-15 um of the midline (the central 30 um).
2. Bilateral vs one-sided. For the remaining components, bilaterality was measured on the magnitude map: the absolute value was taken before binning, and the mirror-cosine between the binned magnitude map and its left-right flip was computed. Its significance was assessed against a random-position null, in which the non-zero voxels were scattered to random positions (∼60 permutations) and the mirror-cosine recomputed, giving a null mean and standard deviation and a z-score z = (observed – null mean)/null SD. A component was bilateral if z >= 2.5, otherwise one-sided.
3. Symmetric vs antisymmetric. Among bilateral components, the sign of the signed binned mirror-correlation (between the signed map and its left-right flip) separated bilaterally symmetric (>= 0; same-sign left/right) from bilaterally antisymmetric (< 0; opposite-sign left/right).

#### Mirrored pairs (functional only)

Because a bilateral left/right pair is often split by ICA into two separate lateralized ensembles, each one-sided functional ensemble was further tested for a mirror partner – a different ensemble in the same recording whose map matches the left-right flip of the one-sided ensemble. A candidate had to be significant both spatially and temporally: the magnitude cross-cosine between one ensemble and the flip of the other exceeded z = 2.5 against the same random-position null (evaluated for all ensemble pairs in a recording as a single matrix product, using each ensemble’s null sigma), and the two calcium time series were significant against a circular time-shift null (|z| >= 2.5), i.e. significantly co-active or anti-active. The best-matching gated partner was kept, and the ensemble was labeled a symmetric pair, split into in-phase (+, time-series correlation r > 0) and anti-phase (–, r < 0). Expression ensembles were classified without a mirrored-partner search (a cross-gene mirror partner is not meaningful).

#### Gene-expression clusters

Gene-expression clusters, from the registered molecular reference atlas decomposition (the same one used for the functional-expression matching in **Fig. 3**), were classified by the identical procedure at 24 um with the expression noise threshold above and no partner pass.

#### Category-average depth maps

For each category in **Fig. 4B**, the per-category mean of the ensemble volumes was computed (each ensemble clipped to its non-negative part before averaging). This mean map was resolved by depth: the reference-frame z-rangFor each category in Fig. 4B, the per-category mean of the ensemble volumes was computed (each ensemble clipped to its non-negative part before averaging). The mirrored-pair maps average only pairs with |r| > 0.3 (2,640 co-active, 856 anti-active). Each mean map was resolved by depth: the reference-frame z-range (excluding the extreme ∼16% ventral and dorsal) was divided into three slabs, and each slab is the maximum over depth of the mean map, with each depth plane normalized to its 99.8th percentile and the same intensity scaling applied to every category. Two categories are overlaid per row by additive color (magenta and green), so voxels present in both appear white.e (excluding the extreme ∼16% ventral and dorsal) was divided into four slabs, and each slab is the maximum over depth of the mean map, with each depth plane normalized to its 99.8th percentile and the same intensity scaling applied to every category.

#### Mediolateral distribution

For each category, we quantified how far its activity mass sits from the midline. Each category’s mean map (above) was reduced to a mediolateral profile, *p*(*x*) = ∑*_y_*_,*z*_ |*v*(*x*, *y*, *z*)|, by summing the absolute map over the anteroposterior (*y*) and dorsoventral (*z*) axes, and the mass-weighted mean absolute distance from the midline was computed as 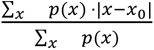, where *x* is the mediolateral center of the reference frame and distances are converted to microns at 3 µm per voxel.

#### Statistics

Our data consist of n = 20,060 functional ensembles, pooled across 68 recordings from 50 larvae (H2B 39 recordings / 32 fish; cyto 29 recordings / 18 fish); recordings lacking a noise-normalized volume were excluded. Category counts (percent of 20,060): bilaterally symmetric 7,494 (37%); mirrored pair 7,977 (40%; co-active 5,424, anti-active 2,553); bilaterally antisymmetric 2,668 (13%); one-sided asymmetric 1,153 (6%); midline 768 (4%). Gene-expression clusters: n = 658 (88 diffuse-noise excluded of 746); symmetric 633 (96%), antisymmetric 14 (2%), midline 11 (2%), one-sided 0. Each component is assigned one category by the ordered rules; three |z| ≥ 2.5 thresholds (bilaterality, pair spatial match, pair time-series correlation) are the only tests.

#### Behavioral regressors

All behavioral variables were tracked from the high-speed behavior camera and processed at their native acquisition rate (tail typically ∼178 Hz) before being placed on each recording’s imaging clock. The non-tail effectors (eyes, mouth, fins) were measured by an unsupervised image decomposition of the relevant video crop: each modality’s cropped movie was read frame by frame, Gaussian-smoothed, downsized, and reduced by incremental PCA, and a small number of dominant components were turned into a one-dimensional motion signal. The channels that entered the models are described below.

● **Eyes.** Eye kinematics came from PCA on the eye videos. The left and right eye crops were combined into a single clip, each frame Gaussian-smoothed (σ = 0.8) and downsized (longest side ≤ 160 px), and reduced by incremental PCA (10 components). The dominant component captures conjugate (yoked) horizontal gaze—the eye position signal—together with a total eye-movement amplitude from the top three components. From this position signal we derived eye *velocity* (its temporal derivative) and an unsigned eye-movement magnitude (|velocity|); the three entered the model as eye position, eye velocity, and eye-movement magnitude. Where separate left/right eye movies existed—or a combined movie could be split at its midline gap— each eye was additionally decomposed on its own: single-eye PC1 is the signed eye-edge rotation (∼66–82% of variance), and a 2-D PCA of the two eyes’ z-scored rotations recovers the conjugate mode on its major axis (identical to the combined regressor) and a convergence/vergence signal on its orthogonal minor axis, plus monocular regressors (each eye’s velocity measured only while the other eye is still). The eye analyses in **Fig. 5** used the conjugate gaze position and its velocity; the convergence and monocular regressors were computed but are not used in this figure.
● **Mouth.** Mouth movement was the incremental-PCA decomposition of the mouth video crop (10 components, per-frame centered, σ = 0.8, ≤ 160 px). The mouth regressor was the absolute temporal derivative, |d/dt|, of the selected component score. The mouth was tracked in only a subset of recordings.
● **Pectoral fins.** Each fin was decomposed separately by incremental PCA (20 components, σ = 1.0, ≤ 220 px). The eigenimages were scored for a left/right dipole structure; the six most dipole-like components were selected, and a fin-amplitude signal was formed as the root-sum-square of those component scores and smoothed over 30 ms. The fin regressor was the absolute temporal derivative of this amplitude, |d/dt|, kept as separate left and right channels. Fins were tracked in only a subset of recordings.
● **Tail deflection and speed.** The native tail-deflection trace (preferring the background-subtracted retracking where available) had a moving-median baseline subtracted, and deflections whose magnitude fell below a MAD-based amplitude threshold were set to zero. The pruned deflection was split into rectified *right* = max(deflection, 0) and *left* = max(−deflection, 0) channels, each also reduced to a one-peak-per-beat binary; *tail speed* was the absolute time derivative, |d/dt(deflection)|. Struggle intervals (below) were zeroed out of all tail-deflection and tail-speed channels over a surrounding window (−1 s to 2 s).
● **Struggle.** Struggle was a computed binary mask: it marked intervals bounded by a matched large left-side (deflection < −threshold) and large right-side (deflection > threshold) event separated by less than 1s (rapid alternating beats), intersected with the detected tail-bout mask. The threshold value was the minimum of 8 degrees and 300 x MAD of the baseline noise.
● **Turn.** Two tail-derived turn signals were available. The grouped-ΔR² turn coupling used a signed net-deflection turn regressor. Turn *direction* for the swim-onset ipsi/contra split and for the exclusive-time left/right-turn analyses used the swim-residualized rectified deflections: the positive part of each side’s deflection after regressing the shared swim drive out of it, so the symmetric swim component is removed, and the signed left−right difference reports direction.
● **Visual stimulus.** For optomotor recordings, left- and right-moving whole-field motion were encoded as separate indicator regressors.

Channels were placed on each recording’s imaging timestamps as follows: the continuous tail-deflection and tail-speed channels were linearly interpolated to the imaging clock, binary bout channels (the tail left/right and struggle binaries) were placed with a binary interpolation, and the fin- , eye-, and mouth-derived channels were anti-alias-averaged onto the imaging clock (a bin-average). Every regressor was then convolved with a jGCaMP8s calcium response kernel. The kernel was a double exponential, *k*(*t*) =*exp*^,*t*/0^*^decay^*−*exp* (−*t*/*τ_rise_*) over a 20 s support, with reporter-specific time constants: *τ_rise_* = 0.03 *s* and *τ_decay_* = 2.0 *s* for H2B-jGCaMP8s (nuclear), and *τ_rise_* = 0.02 *s* and *τ_decay_* = 0.15 *s* for cytosolic jGCaMP8s. Imaging rate varied across recordings, so each regressor was resampled to that recording’s own timestamps. Because not every channel was tracked in every recording, each model used the regressors present for that recording, and any analysis that required a particular channel (e.g., fin or mouth) used only the recordings that contained it.

#### Variable-specific coupling by grouped ΔR^2^

Variable-specific coupling in **Fig. 5C** was quantified with joint ridge-regression models (L2 penalty *α* = 1.0) fitted separately to each ensemble. Related columns were removed as a group: the eye group comprised eye position, eye velocity, and the unsigned eye-movement magnitude; the turn group was the signed net-deflection turn regressor; the swim group was the swim regressor; the struggle group was the struggle regressor; and the visual-stimulus group comprised the left and right stimulus indicators. The grouped ΔR² for a variable was the in-sample R² of the full model minus the R² after refitting with the complete group removed. To estimate the in-sample fitting bias, the ensemble time series was circularly shifted by at least 15 s relative to the fixed design matrix; the full and group-dropped models were refitted and grouped ΔR² recomputed for each of 20 shifts, and the reported effect size was the observed grouped ΔR² minus the mean shift-null value. Clusters represented by fewer than four ensembles were omitted.

#### Stimulus and turn significance across ensembles

For the ranked cluster comparisons (visual-stimulus ipsi/contra in **Fig. 5A** and turn coupling in **Fig. 5C**, left), each ensemble’s value was first averaged within its fish and the fish values were tested; p-values were corrected across all tested clusters with the Benjamini–Hochberg procedure, and clusters were reported as significant at q < 0.05. For turn coupling, the fish means of the null-corrected grouped turn ΔR² were tested against zero with a one-sided Wilcoxon signed-rank test (clusters with at least eight ensembles from at least four fish). For the visual stimulus, each ensemble contributed its mean response during motion toward the ipsilateral side of its hemisphere minus its mean response during motion toward the contralateral side, over the first 80% of the motion window; these differences were averaged within each fish and tested against zero with a one-sample t-test across fish (at least four fish), separately for each reporter line, with the correction applied across all cluster–line pairs.

Stimulus-aligned traces (**Fig. 5A**) were built by aligning each ensemble to the left/right motion epochs, time-normalizing each epoch from motion onset to motion offset onto a common grid, and subtracting a pre-onset baseline; in trials that ended on a swim bout, the last 20% of the epoch was excluded; absolute stimulus laterality is unanchored, so the ipsi/contra split is a relative, within-hemisphere assignment that is internally consistent within each hemisphere convention.

#### Eye-movement coding and population coupling

Three ensembles illustrate distinct eye-movement codes (**Fig. 5B**): *Abd* (eye position), *Abd-Occ* (eye velocity, the temporal derivative of position), and *TriDM* (a slow, smoothed eye position signal). For the example traces, the left and right example ensembles of each cluster were shown over the matched eye regressor (position, velocity, or a low-pass-smoothed position), all drawn from a single recording that carried all three ensembles; ensemble sign was preserved. For the population panel (right), each ensemble was correlated with its ensemble’s target eye signal over the whole recording, and the absolute value of the correlation was used, giving a per-ensemble coupling distribution shown as a boxplot per ensemble. A per-ensemble chance level was obtained by circularly shifting the ensemble by at least 15 s (10 offsets) and recomputing |r| over the same target; the median of this shift-null is drawn as a dashed line. Each ensemble’s observed |r| was compared with its shift-null across ensembles with a paired Wilcoxon signed-rank test across fish, and the three p-values were corrected together with the Benjamini–Hochberg FDR.

#### Turn timescale and derivative coupling

To distinguish fast from slowly integrated turn coupling (**Fig. 5C**), each turn-ensemble was correlated with the low-pass-filtered signed turn regressor across a Butterworth cutoff ladder, folded to the ipsilateral direction by ensemble hemisphere. The same set of ensembles was used at every smoothing window. The curves show the mean across fish. For each cluster, the change in correlation from the native signal to the ∼25 s window was averaged within each fish and tested with a Wilcoxon signed-rank test across fish, and the two p-values were FDR-corrected together. The example panel shows left and right ensembles of *mafaab* and HBO over the signed turn regressor from one recording chosen for well-tracked, discrete turns; the display window is the interval of densest left/right switching, and ensemble sign is preserved. Derivative coupling was illustrated for HDarea by comparing ensemble activity, its temporal derivative, and tail deflection. For the population, each ensemble’s correlation between |d/dt| and the summed left+right turn regressor was averaged within each fish and tested against zero with a one-sided Wilcoxon signed-rank test across fish; the distribution is shown against a circular-shift null.

#### Fin and mouth coupling during behaviorally isolated movements

Because fin and mouth movements were sparse, strongly correlated with each other, and strongly coupled to swimming (**Fig. S5B**), coupling was measured with an exclusive-time absolute correlation. For each ensemble, activity was correlated with the fin or mouth signal only over frames in which that movement was active (z > 1.0) and the tail was quiet (z < 0.5), requiring at least 20 such frames; absolute correlation values were compared with those obtained from 200 random circular shifts of at least 15 s. Fin and mouth were combined as the larger of the available effector-specific correlations to obtain a single orofacial coupling measure. Each ensemble’s value was averaged within its fish; clusters (at least eight ensembles from at least four fish) were ranked by the median of their fish values, and every cluster was tested against the chance level with a one-sided Wilcoxon signed-rank test across fish, with Benjamini–Hochberg correction across all tested clusters.

#### Bout coupling versus orofacial coupling

To ask whether any ensemble is coupled to swim bouts but stays silent during fin/mouth movements (**Fig. 5E**), each ensemble was assessed by two coupling measures that use the data each behavior actually has. The y-axis, *bout coupling*, is the swim-exclusive absolute correlation computed over all recordings: because fin and mouth are tracked in only a minority of recordings, the swim axis uses every recording and reports swim coupling on its own. The x-axis, *orofacial coupling*, is the larger of the fin- and mouth-exclusive absolute correlations, which by construction is measured only when the tail (and the other tracked behaviors) are quiet—that is, it asks whether the ensemble is flat or active during fin/mouth movements that occur without swim. Only recordings with fin or mouth tracking contribute to the x-axis. A cluster above the identity line responds more during swimming than during these fin/mouth-only movements; the five clusters with the largest excess are highlighted. A unique-variance (grouped ΔR²) measure would require fin and mouth in the same model as swim, which most swim recordings lack, so the exclusive-time correlation was used here instead. Clusters were summarized by the median exclusive |r| across their ensembles (at least eight ensembles from at least four fish; exclusive-time active/quiet thresholds and null as in **Fig. 5D**).

#### Peri-struggle-onset dynamics

Struggle bouts were detected with the bout-category struggle detection and each was inspected manually; only struggles judged correct were kept, and struggles resembling escapes, turns or mis-timed detections were discarded. Each ensemble’s activity was z-scored over the recording, resampled around each struggle onset onto a 10-Hz grid from −3 to +7 s, baseline-corrected by the mean over −3 to −1 s, and averaged across struggles within the ensemble, then within fish. Traces show the mean ± s.e.m. across fish. Normal swim bouts (turns and straight swims starting at least 1 s after the previous bout ended) were processed identically. Clusters with at least four ensembles from at least three fish were ranked by their mean response over 0–1.5 s; the top seven and *Vagus* are shown. For each shown cluster, two tests were performed across fish: the rise before struggle onset (mean over −1.0 to −0.2 s minus mean over −3.0 to −1.5 s) compared with the same rise before normal swim bouts (one-sided Wilcoxon signed-rank test on the per-fish differences), and the sustained response (mean over 3–7 s, one-sided Wilcoxon signed-rank test against zero). All 16 p-values were Holm-corrected together. Responses were pooled across struggle directions; ipsiversive and contraversive struggles, classified by net bend relative to the ensemble’s hemisphere, did not differ for any cluster (fish-level Wilcoxon, Holm-corrected).

#### Swim detection, onset latency and lateralized responses

In **Figs. 5G** and **S5C**, swim bouts were detected from native tail speed (typically ∼178 Hz) using a hysteresis threshold on a robust baseline (median + k·MAD): a bout onset was marked when the raw speed rose to at least median + 4·MAD, after having dropped below median + 1.5·MAD (hysteresis re-arm), with a 0.7 s refractory gap between consecutive onsets and a requirement that at least 60% of the 0.5 s preceding an onset lay below the high threshold. Bout onset was set to the start of tail movement: the last time before the bout’s peak at which a combined measure of tail bend and tail speed was at its resting level (median + 1 MAD over the recording). Bouts that began while the tail was already moving were discarded, as were bouts starting less than 1 s after the previous bout ended. Each ensemble time series was z-scored over the recording, resampled around each onset onto a fixed 10-Hz grid, and baseline-corrected by the mean during the preceding second; responses were averaged across bouts within the ensemble. Recordings below 4.5 Hz imaging rate or lacking usable native tail tracking were excluded. For the latency landscape (**Fig. 5G**), a cluster was included when its onset response differed from its pre-onset activity (mean over 0 to +3 s minus mean over −3 to 0 s, two-sided Wilcoxon signed-rank test across ensembles of both reporter lines, Benjamini– Hochberg q < 0.05). Clusters were further required to contain at least eight ensembles from at least four fish and an absolute onset-aligned peak ≥ 0.15; response latency was the first linearly interpolated crossing of half the maximum absolute response, with a 68% bootstrap interval over ensembles (400 resamples; clusters with intervals wider than 0.75 s omitted), and cytosolic GCaMP8s and H2B-GCaMP8s recordings were analyzed separately because of their different kinetics.

For lateralized turn-related ensembles (**Fig. S5C**), bouts were divided into ipsiversive, contraversive, and straight categories. Ensemble hemisphere was determined from the registered spatial map after correcting the left–right mirror introduced by the feature-extraction projection, and each bout was classified by its net bend, the mean tail deflection over its first second minus the median over the 0.2 s before it: a turn if the net bend was at least 2°, with side given by its sign, otherwise straight. A turn was ipsi when the ensemble hemisphere matched the turn direction. An ensemble required at least three bouts in a category, and a category was displayed only when at least eight ensembles and four fish contributed. An ensemble was shown with the directional (ipsi/contra/straight) split only when its ipsiversive and contraversive onset responses differed at p < 0.05 in a paired sign-flip cluster-mass permutation test across fish (cluster-forming threshold 0.05, 2000 permutations); otherwise, ipsiversive, contraversive and straight bouts were merged into one trace per reporter line. Three recordings with unreliable left/right tail tracking (two with a tilted tail, one with all net bends to one side) were excluded from the turn ipsi/contra analysis.

#### Total stimulus- and behavior-explained variance

Total explained variance (**Fig. 6B**) was estimated with a separate lagged ridge model (L2 penalty α = 10). For each recording, the model included all available stimulus and behavior regressors and time-shifted copies of each regressor at −3, −2, −1.5, −1, −0.5, 0, +0.5, +1, +1.5, +2, and +3 s, and was fitted separately to each ensemble’s activity and to its rate of change; each ensemble was scored by the larger of the two null-corrected R² values. Regressors were convolved with the jGCaMP8s kernel and interpolated to the recording’s imaging timestamps before fitting. The model was fitted and scored on the full recording, so the resulting R² is an in-sample fit used descriptively to rank ensembles; a circular-shift refit gave the in-sample overfitting floor, and cross-validated R² for this lagged design stayed near zero for most ensembles and was excluded from the ranking.

#### Statistics and multiple-comparison correction

Unless stated otherwise, each ensemble’s value was averaged within its fish, tests were performed across fish, n reports the number of fish, and error bands and bars are s.e.m. across fish. Cluster-level significance used Wilcoxon signed-rank tests across fish (one-sided against zero or chance for coupling strength; on paired differences for native versus smoothed and struggle versus normal-bout comparisons) and, for the visual-stimulus ipsi/contra difference, a one-sample t-test across fish. Within each family of cluster-level tests (visual-stimulus ipsi/contra; turn coupling; eye population coupling; turn native-versus-smoothed; orofacial coupling), p-values were corrected with the Benjamini–Hochberg procedure (q < 0.05); the struggle tests were Holm-corrected together (**Fig. 5F**), and the bout-onset inclusion gate (**Fig. 5G**) used ensembles as the unit. Ensemble prevalence, the fraction of recordings containing at least one ensemble of the cluster, was compared between darkness and visual stimulation (VR and OMR recordings pooled; **Fig. S5A**) with a Fisher’s exact test per cluster and Benjamini–Hochberg correction across clusters.

#### Activity dynamics and example traces of weakly explained ensembles

Activity kurtosis was calculated from each z-scored ensemble time series as the Fisher (excess) kurtosis (scipy.stats.kurtosis, bias-corrected; Gaussian = 0) and summarized by the median within each ensemble. A descriptive threshold of excess kurtosis ≥ 3 identified sparse, transient activity. Over the full recording, the displayed neurodNet and IthmiNetUnilat ensembles had excess kurtosis 10.6 and 7.3, and the other four displayed ensembles 0.2–2.2.

The six examples in **Fig. 6C**, from the clusters of **Fig. 6B**, were selected from one H2B-jGCaMP8s recording that contained all six together with tail, pectoral-fin and eye tracking. The recording used the virtual reality scene. Ensemble traces were z-scored and shown over a common 120-s interval. The accompanying rows are the jGCaMP8s-convolved design-matrix signals from the same recording and interval: the swim regressor, summed left and right fin activity, and conjugate eye position. Vertical scales are independent across rows.

#### *SGNm-IRr* comparison with single-neuron reconstructions

The final curated SGNm-IRr cluster contained 194 ensembles from 70 recordings. Its across-fish population map (shown in **Fig. 6B**) is the mean of member-ensemble volumes in the symmetric reference space, rendered with the signed gray/magenta ensemble-average convention (see *Spatial-map display*) as top-down and lateral maximum projections.

Molecular organization (**Fig. 6D**) used the positive-loading focal cores (mean|w| + 4 SD|w|) of the three main WARP larvae, whose 42-gene panel allows per-gene enrichment: high-loading neurons assigned to the ensemble were segmented into subclusters as described in **Transmitter identity of spatial subclusters in WARP**, with at least 30 cells per subcluster; subclusters whose dominant transmitter class was serotonergic formed the inferior-raphe group (97 cells) and all others the anterior-hindbrain group (832 cells). Gene enrichment was computed per group as mean expression relative to the hindbrain average. The matched sparse expression component (**Fig. 3D**) and its most strongly correlated source expression maps were taken from the same expression–function matching pipeline (single best-matching component, identical to **Fig. 3D**). *calb2a* was the most enriched gene in the anterior group (11-fold), the second most enriched in the inferior-raphe group after *tph2* (21-fold), and the source map most strongly correlated with the matched expression component (r = 0.66).

For the single-neuron comparison (**Fig. 6E**), reconstructions were assembled in mapzebrain reference space and shown in dorsal and sagittal views on the mapzebrain secondary-gustatory-nucleus and inferior-raphe masks: a ZExplorer connectome query for reciprocal rhombomere-1– inferior-raphe projections (Rh1 → IR and IR → Rh1), and single-neuron reconstructions of secondary-gustatory-nucleus neurons projecting to the right inferior raphe (SGN → IR-R; (Shainer et al., 2023)) together with *tph2*-positive inferior-raphe neurons (Song et al., 2025).

#### Spatial-map display

Unless explicitly stated otherwise, quantitative analyses were performed on registered three-dimensional volumes before display-only contrast adjustment. Figures show top-down maximum projections, often accompanied by a lateral depth projection. Diffuse background was removed for visualization by subtracting a map-specific percentile floor and normalizing to a high percentile except ensemble-average maps, which use a hard-zero baseline and the signed gray/magenta rendering described under **Ensemble average maps and noise normalization.** Display gamma was 0.75 for ensemble-average maps (**Figs. 2E, 3D, 5, 6B** and **S2**), the **Fig. 2B** example ensembles (normalized to the 99.9th percentile) and the **Fig. 6C** ensemble maps; 0.55 for expression components and the **Fig. 3** overlay maps; and 1 (linear) in **Figs. 1**, **2D, 4** and **6A**. Display parameters did not enter regression or symmetry statistics. A common hindbrain silhouette and midline were reused across figures after validation against atlas orientation. Color palettes were selected for color-vision accessibility.

### QUANTIFICATION AND STATISTICAL ANALYSIS

Analysis-specific sample sizes are reported in the corresponding Methods sections and figure legends. The independent observational unit was the ensemble for ensemble-level regression and bootstrap analyses, the recording for behavior–behavior cross-correlograms, and the fish for the all-ensemble peri-swim-onset supplement. Where multiple ensembles from one fish were displayed or summarized, this nesting is stated explicitly.

Boxplots show the median, interquartile range and 1.5× interquartile-range whiskers. Traces show mean ± s.e.m. across the observational units specified for the analysis. Variable-specific grouped ΔR² distributions were assessed with one-sided Wilcoxon signed-rank tests against zero after shift-null correction, with α=0.05; the separate median ≥0.05 rule was an effect-size support criterion. Where a single analysis tested many ensembles in parallel and used that test to select or rank the ensembles shown, the per-ensemble p-values were corrected for multiple comparisons with the Benjamini– Hochberg false-discovery-rate procedure at q = 0.05: the turn-coupling population ranking, the ipsiversive-versus-contraversive stimulus-response split, the swim-onset response gate used for the response-timing panel, and the stimulus-prevalence contrasts (a per-ensemble Fisher’s exact test). The per-ensemble grouped-ΔR² significance calls used to flag individual behavior-panel ensembles (one-sided Wilcoxon signed-rank with the median ≥ 0.05 support rule, as above) were reported without multiple-comparison correction.

Latency uncertainty was estimated from 400 bootstrap resamples across ensembles and reported as the 16th–84th percentile interval. Pearson correlation was used for spatial cosine-label validation, mirror-symmetry scale curves, partner time-series association and behavior cross-correlation as described above. Symmetry categories and recovery calls were deterministic threshold- or curation-based classifications and were not subjected to inferential tests.

Analyses were implemented in Python 3.11.7 using NumPy 1.26.4, SciPy 1.17.1, scikit-learn 1.8.0, scikit-image 0.22.0, nibabel 5.2.0, and Matplotlib 3.10.9, on a workstation running Windows 11 with a 16-core AMD processor and 128 GB of RAM.

#### AI-assisted code development

OpenAI Codex GPT 5.4 (OpenAI) and Claude Code Opus 4.8 (Anthropic) were used as programming assistants to support the writing, refactoring, and debugging of portions of the custom research code. These tools were not used as analytical models on the experimental data. Analytical objectives and methodological choices were specified by the authors, and all AI-assisted code was reviewed, tested, and modified as needed before use. The authors examined the resulting analyses and outputs and take responsibility for the validity and reproducibility of the final code and results.

## SUPPLEMENTAL INFORMATION

**Supplementary Figure S1.**
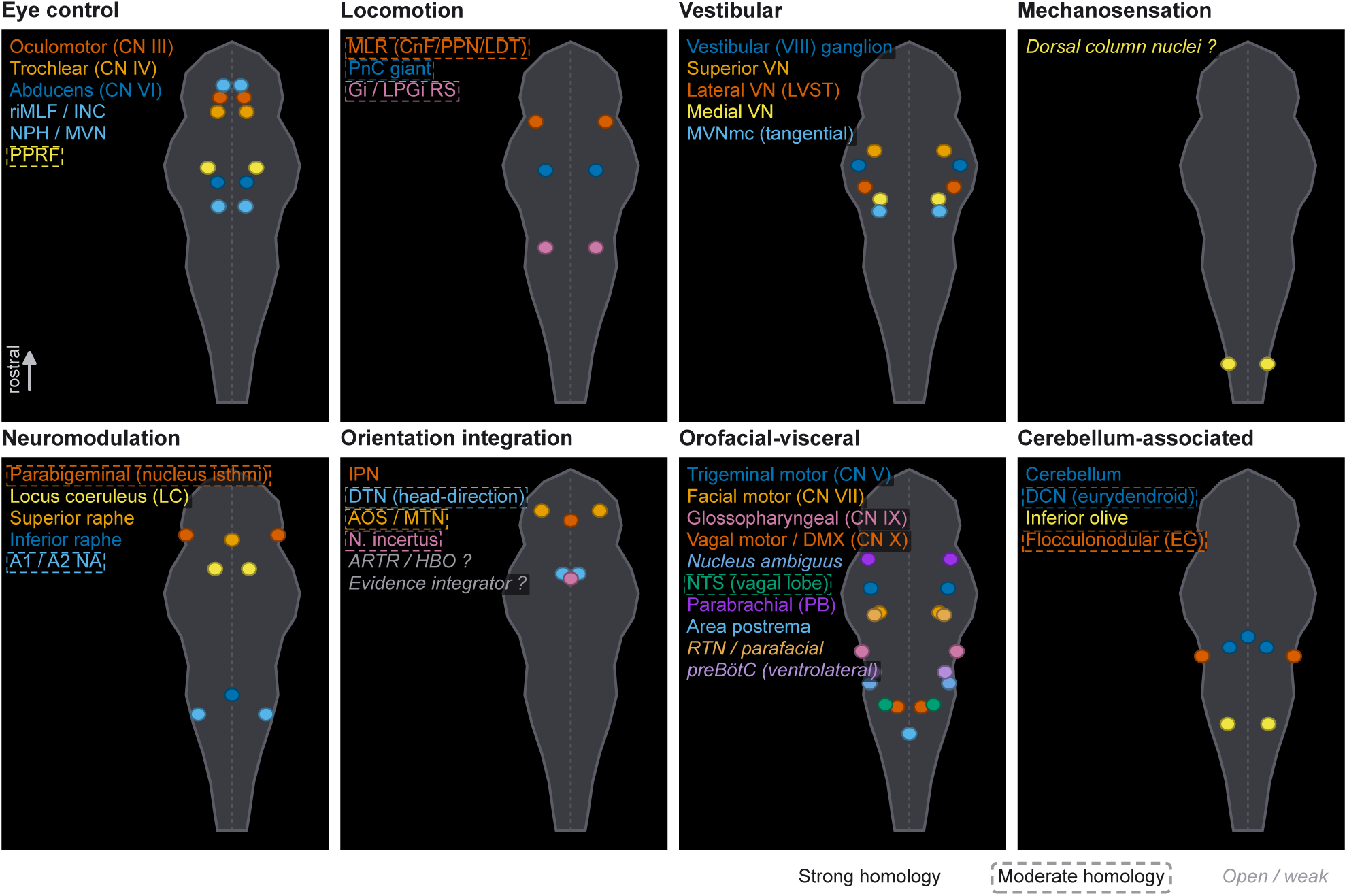
Proposed mammalian counterparts of established zebrafish hindbrain populations, illustrated schematically in the mouse. Proposed mouse counterparts of the zebrafish populations in Fig. 1 are shown as colored dots on a schematic brainstem background (gray; rostral at the top). The diagram indicates approximate positions only; dot size and spacing do not represent anatomical size or spatial relationships. Panels follow the same eight functional groups, with colors matching the corresponding zebrafish populations. Homology confidence is indicated by label formatting: strong correspondences have unboxed regular labels, moderate correspondences have dashed boxes, and open/weak correspondences have italic labels. Open/weak correspondences indicate unresolved or candidate relationships, including zebrafish populations for which no specific mammalian counterpart has yet been established. **Supplementary Table 1** details the supporting evidence, caveats, and references for each assignment. **Abbreviations: Eye control:** NPH, nucleus prepositus hypoglossi; PPRF, paramedian pontine reticular formation; riMLF, rostral interstitial nucleus of the medial longitudinal fasciculus. **Locomotion:** CnF, cuneiform nucleus; Gi, gigantocellular reticular nucleus; INC, interstitial nucleus of Cajal; LDT, laterodorsal tegmental nucleus; LPGi, lateral paragigantocellular nucleus; MLR, mesencephalic locomotor region; PnC, caudal pontine reticular nucleus; PPN, pedunculopontine nucleus; RS, reticulospinal. **Vestibular:** LVST, lateral vestibulospinal tract; MVN, medial vestibular nucleus; MVNmc, magnocellular subdivision of the medial vestibular nucleus; VN, vestibular nucleus. **Neuromodulation:** A1/A2, medullary noradrenergic cell groups; LC, locus coeruleus; NA, noradrenergic/catecholaminergic groups. **Orientation integration:** AOS, accessory optic system; ARTR, anterior rhombencephalic turning region; DTN, dorsal tegmental nucleus; HBO, hindbrain oscillator; IPN, interpeduncular nucleus; MTN, medial terminal nucleus. **Orofacial–visceral:** DMX, dorsal motor nucleus of the vagus; NTS, nucleus of the solitary tract; preBötC, preBötzinger complex; RTN, retrotrapezoid nucleus. **Cerebellum-associated:** DCN, deep cerebellar nuclei; EG, eminentia granularis. **General:** CN, cranial nerve; N., nucleus.

**Supplementary Figure S2.**
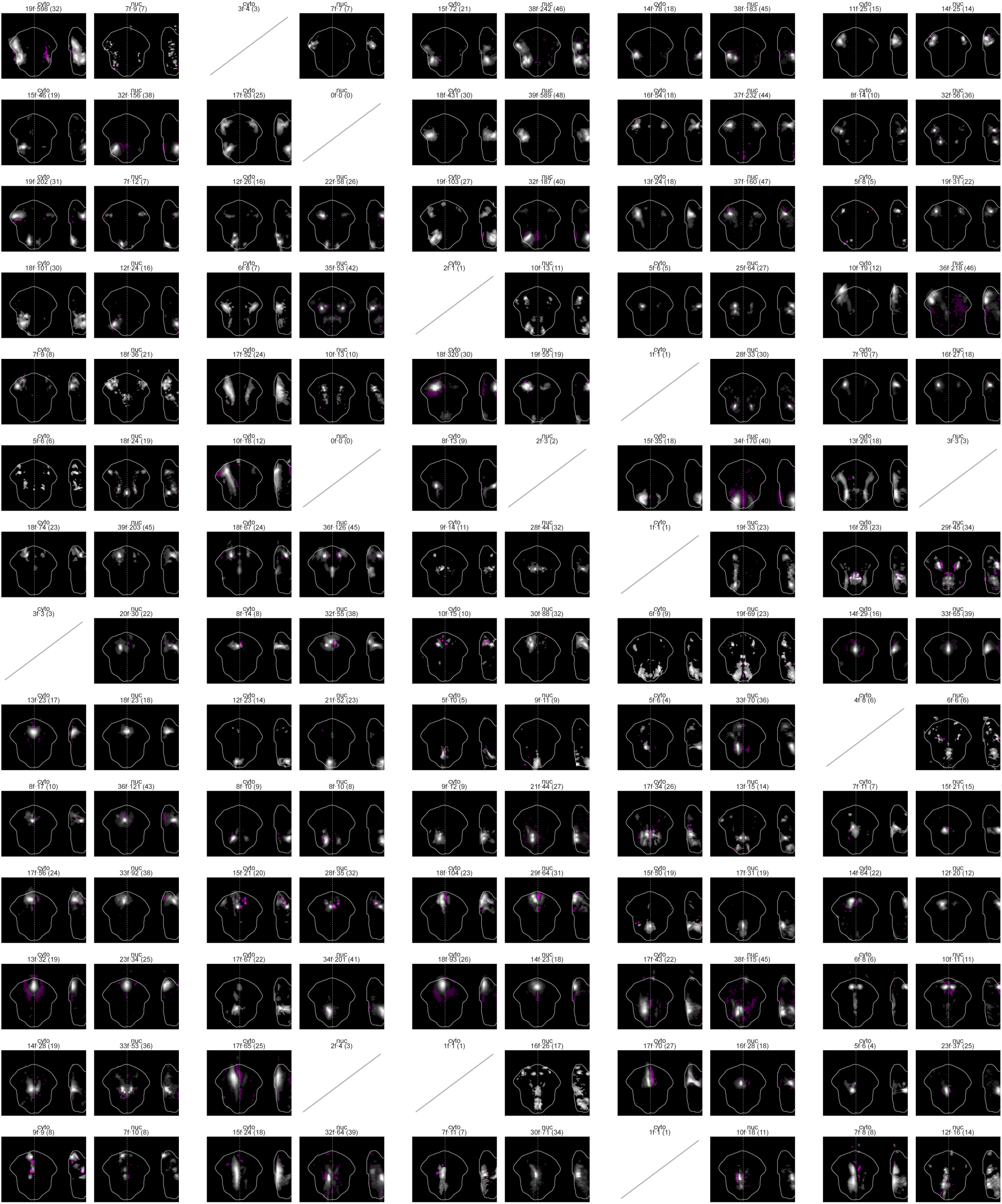
Curated ensemble average maps are consistent across the two reporter lines. Each curated ensemble from Fig. 2E is shown separately for the cytosolic jGCaMP8s (“cyto,” left) and nuclear-localized H2B-jGCaMP8s (“nuc,” right) lines. A map is shown only when at least five fish from that reporter line contributed to the ensemble; otherwise, the corresponding position is marked by a diagonal line. For each reporter line, ensemble maps were noise-normalized, thresholded, and oriented with their stronger hemisphere on the left before averaging. Average maps are shown as signed maximum-intensity z-projections (gray, positive weights; magenta, negative weights), with the hindbrain outline and midline overlaid, and a YZ depth projection at right. Voxels were retained where the mean exceeded 4/√n in noise units (n = ensembles from that line); maps use a hard-zero baseline, per-map normalization to the 99th percentile of projected magnitude and a display gamma of 0.75. Labels give the numbers of contributing fish, ensembles, and recordings for each reporter line.

**Supplementary Figure S3.**
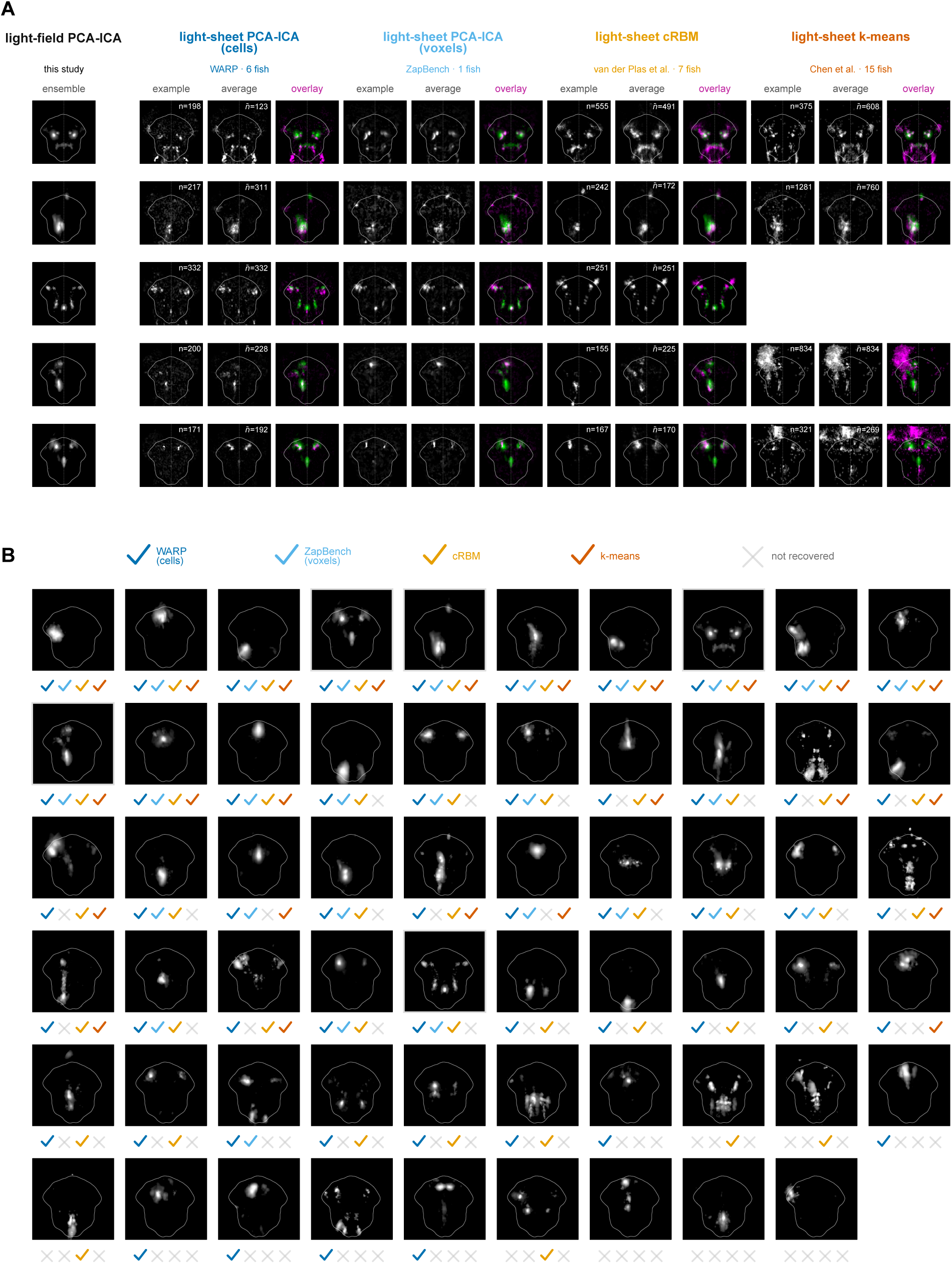
Hindbrain ensembles are recovered in independent datasets using alternative analysis methods. Ensembles from four publicly available whole-brain light-sheet datasets were registered to the common light-field reference space (150 × 150 × 76 voxels) and matched to our ensemble averages using a common spatial-similarity measure, followed by manual curation (**Methods**). The four analysis methods were spatial PCA–ICA applied to segmented cells from WARP recordings (Marquez-Legorreta et al., 2026); 3 fish), spatial PCA–ICA applied voxel-wise to ZapBench data (Lueckmann et al., 2025); 1 fish), compositional restricted Boltzmann machines (cRBM; (Van Der Plas et al., 2023); 7 recordings), and k-means correlation clustering (*k* = 20; (Chen et al., 2018); 15 fish). The two PCA–ICA datasets were treated as separate methods in the recovery analysis. All maps were left–right symmetrized before matching, allowing a lateralized ensemble to match its mirrored counterpart. **A,** Example matched ensembles. The leftmost column shows the positive part of our across-fish light-field ensemble average (Fig. 2E). For each method, the subsequent columns show one matched ensemble, the mean map of all ensembles assigned to that cluster, and an overlay of our average in green with the mean for the different method in magenta. Blank entries indicate that the ensemble was not recovered by that method. For cell-based methods, *n* indicates the number of hindbrain member neurons in the displayed ensemble and *n̄* the mean across matched ensembles. **B,** Recovery across the eligible ensemble set. Of the 70 clusters shown in Fig. 2E, 59 were eligible for cross-dataset comparison. We excluded 11 clusters for which ≥80% of ensembles came from the cytosolic jGCaMP8s line, because the comparison datasets did not reproduce these neuropil-rich cytosolic ensembles. Each thumbnail shows one eligible ensemble average. The four symbols beneath each thumbnail indicate recovery by cell-wise PCA–ICA, voxel-wise PCA–ICA, cRBM, and k-means, respectively. A colored checkmark indicates recovery, defined as at least one ensemble assigned to the cluster; a gray × indicates that no match was identified by that method, but does not imply that the ensemble was absent from the dataset.

**Supplementary Fig. S4.**
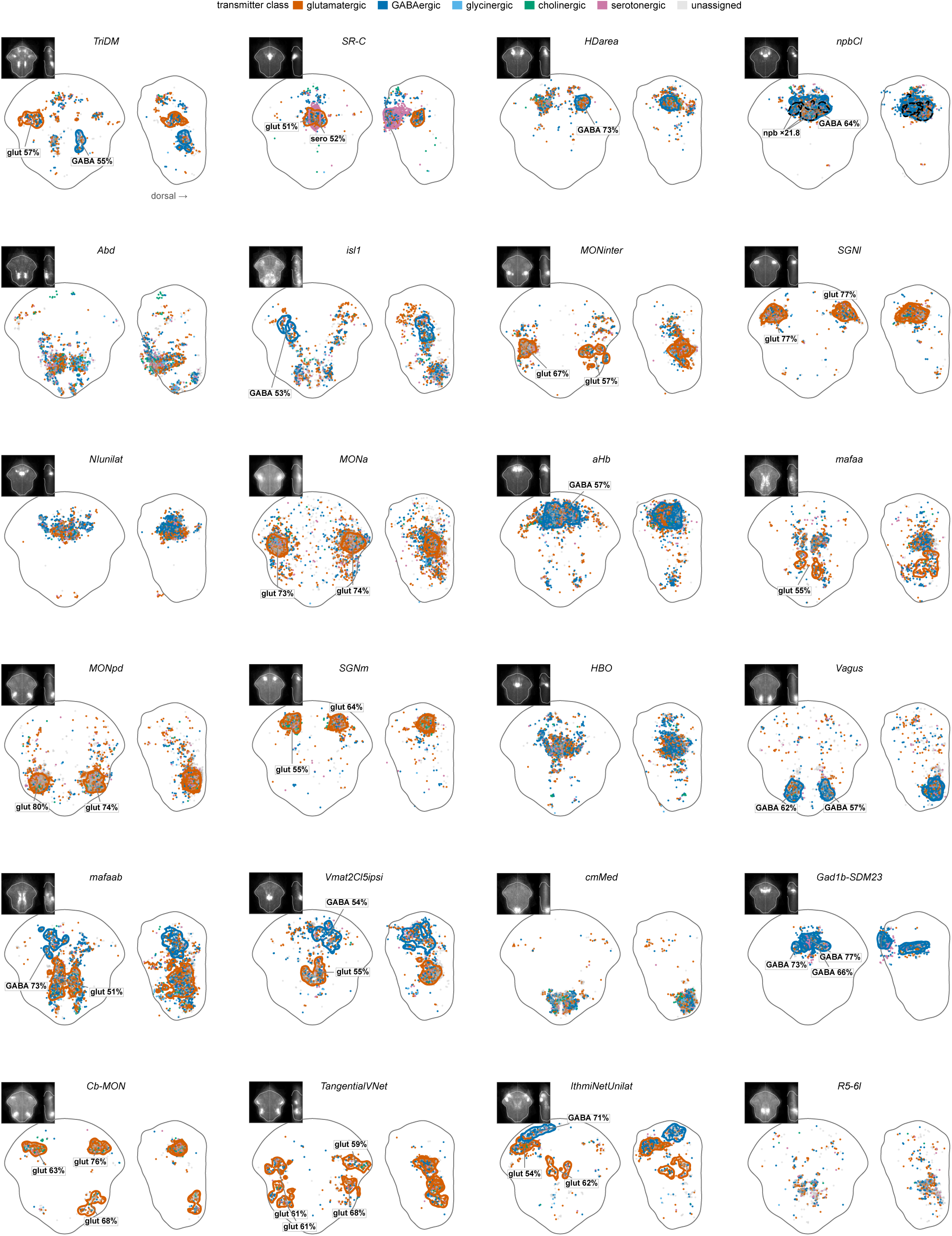
Transmitter identity and internal molecular organization of ensembles recovered in an independent dataset. For 24 functional ensembles (see Table 1 for all ensembles), the high-loading neurons of every ensemble assigned to that cluster in the WARP light-sheet dataset ((Marquez-Legorreta et al., 2026), **Fig. S3**), pooled across up to six larvae, and restricted to the functional hindbrain mask, are colored by transmitter identity measured by multiplexed HCR in the same cells; unassigned neurons are light gray. Left, dorsal view (rostral up); right, depth view at the same scale (dorsal to the right); the gray line outlines the hindbrain within the functional mask, and the inset shows the light-field ensemble average. High-loading neurons were segmented into localized subclusters; a subcluster is outlined and labeled with its dominant transmitter class where that class exceeds 50% of its typed cells and the subcluster holds at least 10% of the ensemble’s typed cells, with percentages computed over typed cells only. *HBO* was divided into a medial and a lateral part at the gap in its neurons’ distance from the midline. Dashed black outlines in *npbCl* mark clusters of npb-positive neurons, labeled with their fold enrichment over the hindbrain mean. Transmitter class was assigned from abundance-normalized marker counts (**Methods**); cholinergic identity comes from the three main larvae only.

**Figure S5.**
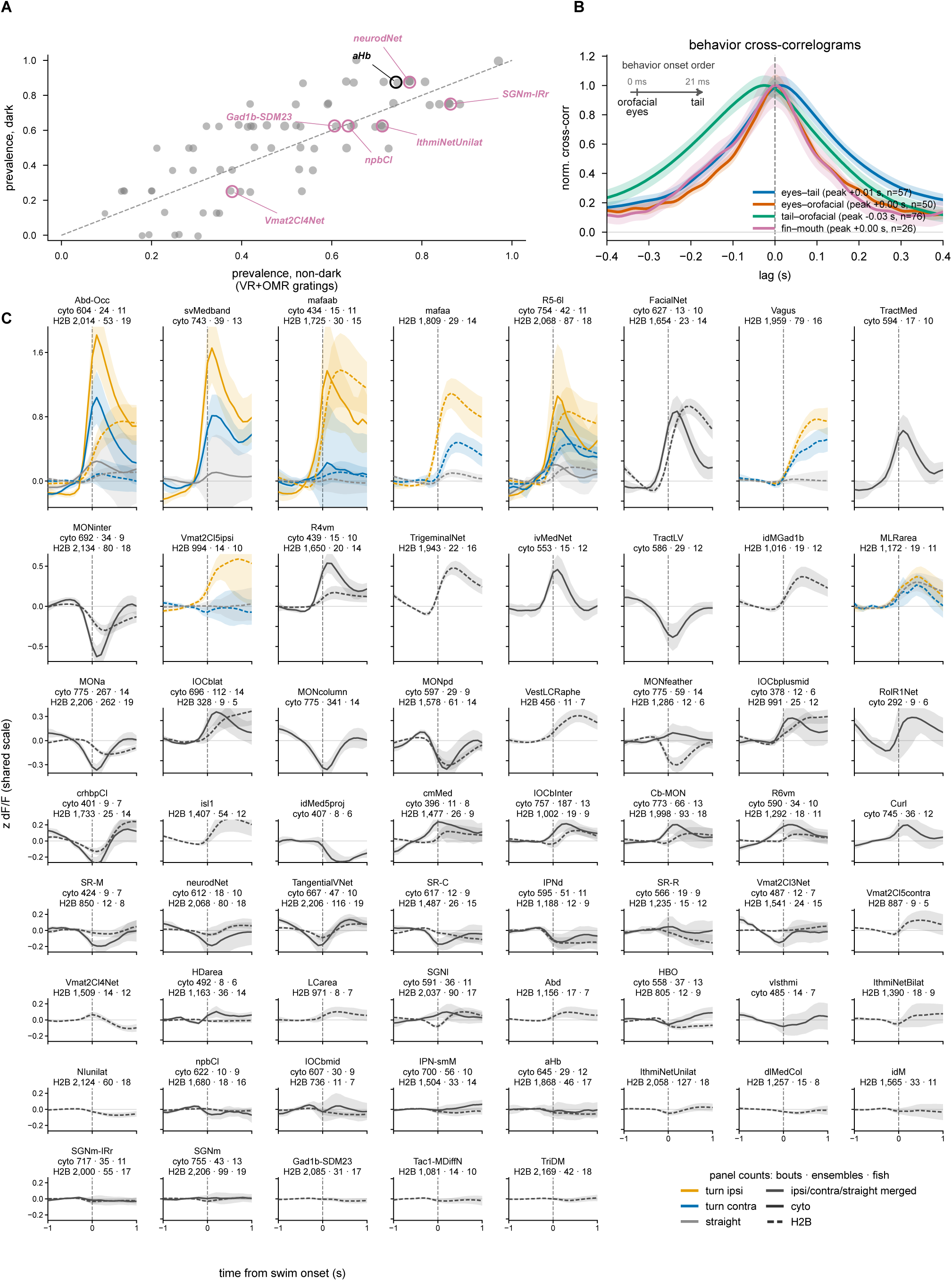
Ensemble occurrence across stimulus conditions, relative timing of tracked behaviors, and swim-onset responses across the ensemble atlas. **A,** Ensemble occurrence does not depend on visual stimulation. For each ensemble, the fraction of recordings in which it was detected is compared between recordings in darkness (y) and recordings with visual stimulation (closed-loop VR + OMR gratings). No ensemble differed significantly (0 of 70; Fisher’s exact test per cluster, Benjamini–Hochberg FDR q < 0.05; 66 recordings with visual stimulation, 8 in darkness). Circles mark aHb (black) and the six ensembles of Fig. 6B (magenta). **B,** Cross-correlograms of eye, tail, and orofacial movement speed (magnitude of the native-rate time-derivative (|d/dt|) of each tracked signal (**Methods**)). Curves show mean ± s.e.m. across recordings, each normalized to its peak; the key gives each pair’s peak lag and number of recordings, and the inset orders behavior onsets by these lags. **C,** Peri-swim-onset responses across the ensemble atlas, mean ± s.e.m. across fish over −1 to +1 s around swim onset; cytosolic-jGCaMP8s solid, H2B-jGCaMP8s dashed lines; panels ordered by response amplitude, largest first, with the top three rows drawn taller at the same dF/F scale. For the 8 of 61 ensembles whose ipsiversive and contraversive responses differed (p < 0.05, uncorrected), turn-ipsi (orange), turn-contra (blue), and straight (gray) bouts are shown separately; for the others, they are merged. Panel titles give the numbers of bouts, ensembles, and fish for each reporter line whose curve is shown.

**Supplementary Table 1.** Established zebrafish hindbrain functional populations: functions, mask provenance, and proposed mammalian counterparts.

